# An ESCRT-LEM domain inner nuclear membrane protein surveillance system is poised to directly monitor the integrity of the nuclear envelope barrier and nuclear transport system

**DOI:** 10.1101/523670

**Authors:** David J. Thaller, Matteo Allegretti, Sapan Borah, Paolo Ronchi, Martin Beck, C. Patrick Lusk

**Author notes:** Correspondence to C. Patrick Lusk.

## Abstract

The integrity of the nuclear envelope membranes coupled to the diffusion barrier and selective transport properties of nuclear pore complexes (NPCs) are a prerequisite for the robust segregation of nucleoplasm and cytoplasm. Recent work supports that mechanical membrane disruption or perturbation to NPC assembly can trigger an ESCRT-dependent surveillance system that seals nuclear envelope pores: how these pores are sensed and sealed remains to be fully defined. Here, we show that the principal components of the nuclear envelope surveillance system in yeast, which includes the ESCRT Chm7 and the integral inner nuclear membrane (INM) protein Heh1, are spatially segregated by the nuclear transport system. Specifically, at steady state Chm7 is actively restricted from the nucleus by Crm1/Xpo1. Consistent with the idea that it is the exposure of the INM that triggers surveillance, the expression of a transmembrane anchor and the winged helix domain of Heh1 is sufficient to recruit and activate Chm7 at a membrane interface. Correlative light electron tomography under conditions of Chm7 hyper-activation further show the formation of an elaborate network of fenestrated sheets at the INM and suggest ER-membrane delivery at sites of nuclear envelope herniation. Our data point to a model in which exposure of Chm7 to Heh1, driven by any perturbation in the nuclear envelope barrier would lead to local nuclear envelope remodeling to promote membrane sealing. Our findings have implications for disease mechanisms associated with defects in NPC assembly and nuclear envelope integrity.

## Introduction

The molecular machinery that biochemically segregates the nucleus and the cytoplasm has been extensively investigated. The foundational components of this selective barrier include the double-membrane nuclear envelope with embedded nuclear pore complexes (NPCs). NPCs impose a “soft” diffusion barrier to macromolecules larger than ~40 kD (Popken et al., 2015; Timney et al., 2016) while providing binding sites for the rapid and selective transport of signal-bearing (nuclear localization and nuclear export signals; NLSs and NESs) macromolecules, which are ferried through the NPC by shuttling nuclear transport receptors (NTRs; a.k.a. karyopherins/importins/exportins; (Schmidt and Görlich, 2016)). Directionality and energy for NTR-selective transport is imparted by the spatial segregation of the Ran-GTPase whose nuclear GTP-bound form destabilizes and stabilizes import and export complexes, respectively (Floch et al., 2014).

Interestingly, the robustness of the nuclear envelope barrier has been shown to be compromised in several different contexts, including in diverse human diseases (Hatch and Hetzer, 2014; Lusk and King, 2017). For example, there is an emerging body of work linking the function of NTRs and NPCs with neurodegenerative diseases like ALS and FTD (Nousiainen et al., 2008; Freibaum et al., 2015; Jovičić et al., 2015; Kaneb et al., 2015; Zhang et al., 2015; Kim and Taylor, 2017; Shi et al., 2017). These studies, coupled to the observations of age-related declines in NPC function in both post-mitotic multicellular systems (D’Angelo et al., 2009; Savas et al., 2012; Toyama et al., 2013) and also in replicative aging models like budding yeast (Janssens et al., 2015; Lord et al., 2015), support a theme in which the function of the nuclear envelope could be mitigatory of age-related disease progression (Schreiber and Kennedy, 2013; Jevtić et al., 2014; Serebryannyy and Misteli, 2018). Of similar interest, the hallmark cellular pathophysiology of early-onset dystonia are nuclear envelope herniations (Goodchild et al., 2005) that emanate from NPC-like structures (Laudermilch et al., 2016). As analogous herniations have been observed in many genetic backgrounds associated with defects in NPC biogenesis in yeast over several decades (Thaller and Lusk, 2018), this has contributed to the idea that the herniations might be the result of either defective NPC assembly events (Scarcelli et al., 2007; Onischenko et al., 2017; Zhang et al., 2018) and/or the triggering of a NPC (Wente and Blobel, 1993) or NPC assembly quality control pathway (Webster et al., 2014, 2016). The latter could depend on the function of the endosomal sorting complexes required for transport (ESCRT), a membrane scission machinery that has been proposed to seal-off malforming NPCs (Webster et al., 2016).

That there could be mechanisms to surveil the assembly of NPCs makes considerable sense as there are hundreds of NPCs, each containing hundreds of nucleoporins/nups (Kosinski et al., 2016; Kim et al., 2018) that are assembled during interphase in mammalian cells (Maul et al., 1972; Doucet et al., 2010; Dultz and Ellenberg, 2010). There are approximately 100 NPCs formed during a budding yeast cell cycle, which includes a closed mitosis (Winey et al., 1997). As interphase NPC assembly likely occurs through an inside-out evagination of the inner nuclear membrane (INM) followed by membrane fusion with the outer nuclear membrane (ONM)(Otsuka et al., 2016), holes are constantly being formed in the nuclear envelope. Without mechanisms to surveil this process, *de novo* NPC biogenesis might pose a threat to nuclear-cytoplasmic compartmentalization (Webster et al., 2014). Consistent with this idea, malformed or damaged NPCs are not passed on to daughter cells in budding yeast (Colombi et al., 2013; Makio et al., 2013; Webster et al., 2014) and deletion of the ESCRT machinery in the context of genetic backgrounds where nuclear envelope herniations have been observed e.g. *nup116Δ* (Wente and Blobel, 1993) or *apq12Δ* (Scarcelli et al., 2007) cells require a nuclear envelope-specific ESCRT, Chm7 (the orthologue of mammalian CHMP7) for viability (Bauer et al., 2015; Webster et al., 2016). While we have previously proposed that a biochemical signature of malforming NPCs is surveilled by integral inner nuclear membrane proteins of the LAP2-emerin-MAN1 (LEM) domain family, specifically Heh2, it remains to be formally established what the signal that leads to ESCRT recruitment to the nuclear envelope actually comprises (Webster et al., 2014).

Evidence that the ESCRT machinery acts at holes in the nuclear envelope is further exemplified by their critical role in performing annular fusion events during the final stages of nuclear envelope reformation at the end of mitosis in mammalian cells (Olmos et al., 2015, 2016; Vietri et al., 2015; Gu et al., 2017; Ventimiglia et al., 2018). Moreover, ESCRTs are also required for the efficient repair of nuclear ruptures that arise during the migration of cells through tight constrictions (Denais et al., 2016; Raab et al., 2016). And, it is most likely that they also act to repair nuclear envelope ruptures that are induced by intracellular mechanical stresses from either the actin cytoskeleton (Hatch and Hetzer, 2016; Robijns et al., 2016), or from those observed during telomere crisis (Maciejowski et al., 2015). Lastly, recent work also suggests a role for ESCRTs in the context of turning over NPCs in terminally differentiated cells (Toyama et al., 2018). It remains an open question, however, whether the mechanisms that repair nuclear ruptures, seal the nuclear envelope at the end of mitosis, and protect against defective NPC assembly respond to an identical upstream signal and proceed through the same membrane-sealing mechanism.

Clues to what might constitute the upstream signal that leads to nuclear envelope-recruitment of ESCRTs could be drawn from other contexts where ESCRTs protect membrane compartments including endolysosomes (Skowyra et al., 2018; Radulovic et al., 2018) and the plasma membrane (Jimenez et al., 2014; Scheffer et al., 2014; Gong et al., 2017). In both of these cases, there is evidence to suggest that the local release of Ca^2+^ is a trigger for ESCRT recruitment, through (at least at the plasma membrane) a Ca^2+^ binding protein, ALG-2 (Jimenez et al., 2014; Gong et al., 2017). Whether Ca^2+^ plays a role at the nuclear envelope remains unaddressed. More generally, there are two, often redundant, recruitment mechanisms seeded by either an ESCRT-I, II complex and/or ESCRT-II and ALIX (Bro1 in yeast) that bind and activate ESCRT-III subunit polymerization (Wemmer et al., 2011; Henne et al., 2012; Tang et al., 2015, 2016; Christ et al., 2016) on specific membranes throughout the cell (reviewed in (Schöneberg et al., 2017; McCullough et al., 2018)).

ESCRT-III polymers predominantly made up of the most abundant ESCRT-III (Snf7/CHMP4B) scaffold negative but also in at least one case, positive membrane curvature (McCullough et al., 2015), and directly contribute to membrane scission (Adell et al., 2014, 2017; Schöneberg et al., 2018). The AAA+ ATPase Vps4 disassembles ESCRT-III filaments by directly interacting with MIM (MIT interacting motif) domains present on a subset of ESCRT-III subunits (Obita et al., 2007; Stuchell-Brereton et al., 2007; Agromayor et al., 2009; Xiao et al., 2009; Han et al., 2015) by threading the ESCRT-III filaments through the central cavity of a hexameric ring (Yang et al., 2015; Han et al., 2017; Monroe et al., 2017; Su et al., 2017). It is likely that ESCRT-III disassembly by Vps4 directly contributes force to promote membrane scission (Schöneberg et al., 2018). Whether the membrane scission reaction is different in distinct subcellular contexts like at the nuclear envelope remains to be understood.

Consistent with the idea that there might be unique ESCRT membrane remodeling mechanisms at play in distinct compartments, a step-wise recruitment and activation mechanism requiring the ESCRT-II Vps25 and the ESCRT-III Vps20 is thought to be required at budding yeast endosomes (Saksena et al., 2009; Teis et al., 2010; Tang et al., 2015, 2016), but both of these proteins are absent from the genetic and biochemical analyses of the nuclear envelope arm of the ESCRT pathway (Webster et al., 2014, 2016). These data suggest that other proteins likely contribute to ESCRT-III activation at the nuclear envelope. Key candidates are Chm7 and the inner nuclear membrane (INM) proteins, Heh1/Src1 (orthologue of LEM2) and Heh2 (orthologue of MAN1 or other LEM-domain proteins). These proteins have collectively been shown to interact biochemically and genetically with Snf7 (Webster et al., 2014, 2016) and Heh1 is required for the focal accumulation of Chm7 at the nuclear envelope in genetic backgrounds where NPC assembly is inhibited (Webster et al., 2016). Remarkably, the interactions between Heh1 and Chm7 are well conserved in both fission yeast (Gu et al., 2017) but also in mammalian cells, where LEM2 is required to recruit CHMP7 to the reforming nuclear envelope at the end of mitosis (Gu et al., 2017). It remains to be understood, however, whether LEM proteins (or Chm7) directly contribute to ESCRT-III activation at the nuclear envelope or whether additional proteins are involved.

Heh1 and Heh2 contain an N-terminal helix-extension-helix (heh) motif (the LEM domain), followed by an INM targeting sequence that (at least in the case of Heh2 but likely also Heh1 (King et al., 2006; Lokareddy et al., 2015)) includes an NLS and a ~200 amino acid unstructured region that are both required for INM targeting (Meinema et al., 2011). They both also contain a second nuclear-oriented domain, which likely folds into a winged helix (WH)(also called MAN1/Src1-C-terminal homology domain or MSC; (Caputo et al., 2006)); this domain is also well conserved through evolution (Mans et al., 2004; Mekhail et al., 2008). The LEM domain proteins as a family have been ascribed diverse roles in gene expression either through binding to transcription factors, BAF, or the lamins (Barton et al., 2015). While yeasts lack BAF and lamins, the LEM domain proteins nonetheless directly interface with chromatin (Grund et al., 2008; Barton et al., 2015), contribute to rDNA repeat stability (Mekhail et al., 2008) and the mechanical robustness of the nucleus (Schreiner et al., 2015). This latter function is likely revealed by the observation in many diverse yeast species that loss of Heh1 leads to nuclear envelope disruption (Yewdell et al., 2011; Gonzalez et al., 2012; Yam et al., 2013), however, it might also reflect Heh1’s role in recruiting ESCRTs to the nuclear envelope. Thus, the relationship between how the LEM proteins contribute to nuclear integrity and ESCRT-mediated surveillance remains to be clearly defined.

In the following, we further explore the molecular determinants of Chm7 recruitment to the budding yeast nuclear envelope by Heh1. We determine that the spatial segregation of Chm7 and Heh1 on either side of the nuclear envelope is driven by NTRs and the robustness of the nuclear transport system. Perturbations of this system, or the exposure of Heh1 to the cytosol leads to the local recruitment and Heh1 WH-dependent activation of Chm7. At sites of Chm7 hyperactivation, we observe remarkable alterations to nuclear envelope morphology including nuclear envelope herniations and intranuclear INM invaginations suggesting a role for membrane expansion and remodeling during nuclear envelope repair.

## Results

### Chm7 is actively exported from the nucleus by Xpo1

It was our previous experience that visualizing endogenous levels of Chm7-GFP was challenging due to its low level of expression (Webster et al., 2016), thus, to gain further insight into the localization determinants of Chm7 in budding yeast, we overexpressed Chm7-GFP behind the control of a galactose (*GAL1*) inducible promoter. As shown in Figure 1C, culturing of cells in galactose for a short time (~45 min) led to the appearance of Chm7-GFP fluorescence localized throughout the cytosol. Unexpectedly, we also observed that Chm7-GFP was excluded from the nuclear interior (orange asterisks). These data raised the possibility that Chm7-GFP is unable to cross the diffusion barrier imposed by NPCs, or, there is an active nuclear export pathway that prevents Chm7 from accessing the nucleus.

**Figure 1:**
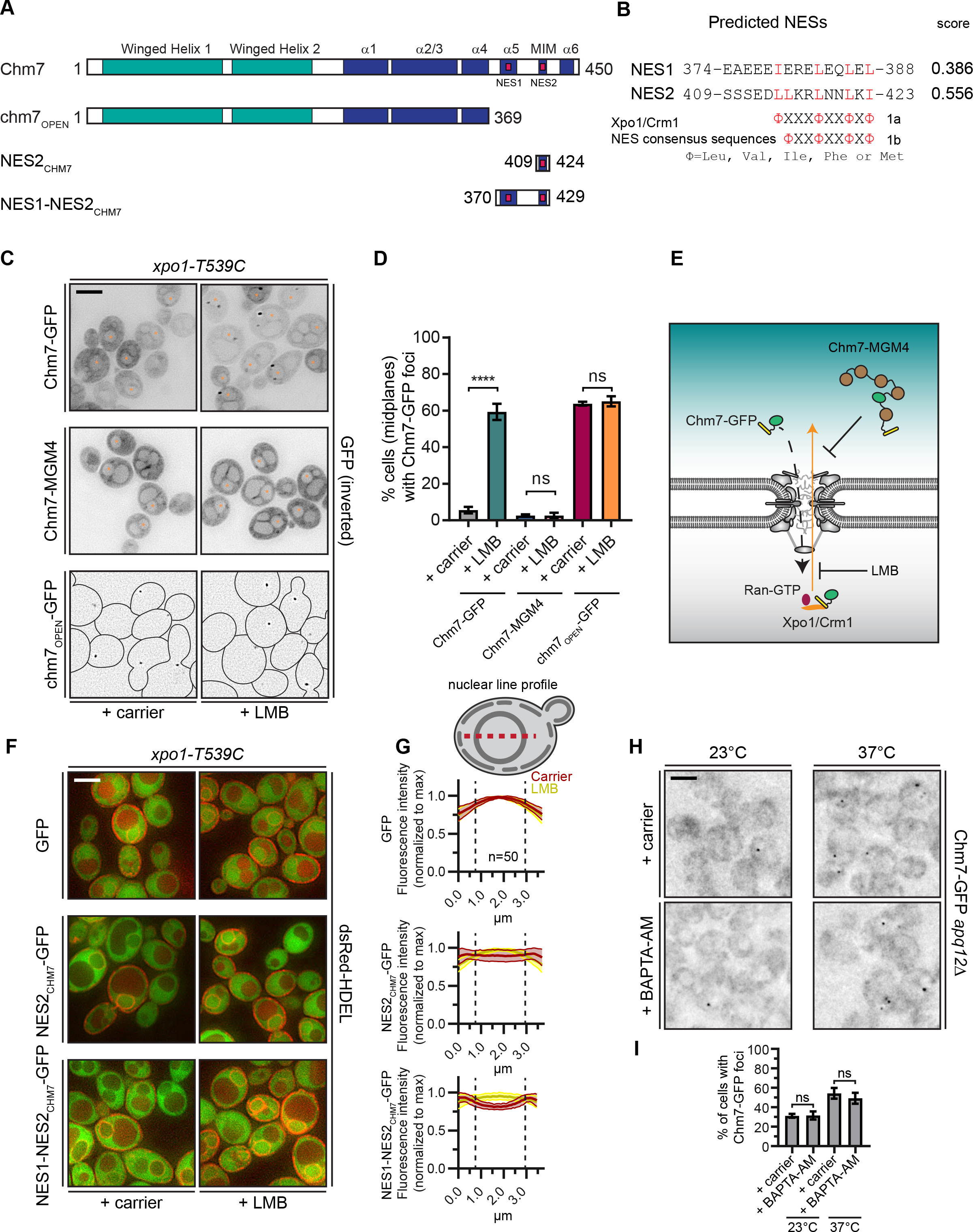
Chm7 can diffuse across the NPC but is actively exported by Xpo1. **A**) Schematic of Chm7 and deletion constructs with predicted winged helix domains (teal), alpha helices (blue) and NESs (red boxes); numbers are amino acids. **B**) Predicted Chm7 NESs with probability score from LocNES; numbers are amino acids from Chm7 sequence. Hydrophobic residues in putative NESs are highlighted red as per the consensus class 1a and 1b NESs shown. **C**) Deconvolved inverted fluorescence micrographs of the indicated overexpressed Chm7-GFP constructs in a LMB-sensitive strain (*xpo1-T539C*) treated with carrier (MeOH) or LMB. Nuclei are marked with orange asterisks. Cells are outlined in black in bottom panels. Scale bar is 5 μm. **D**) Plots showing the percentage of cells with Chm7-GFP nuclear envelope foci from C. Data are from three independent replicates where >100 cells were counted for each strain. Only images of midplanes were quantified. *P*-values from unpaired Student’s t-test where ns is *P* > 0.05, **** *P* ≤ 0.0001. **E**) Schematic of the experiment and interpretation of C. **F**) Deconvolved fluorescence micrographs (merge of green and red channels) of the indicated GFP, and GFP-NES constructs co-expressed with dsRed-HDEL to help visualize the nucleus. Cells were treated with carrier (MeOH) or LMB before imaging. Scale bar is 5 μm. **G**) To quantify relative nuclear exclusion of the GFP and GFP-NES constructs in F, line profiles bisecting the nucleus as shown in diagram were measured from 50 cells/condition pooled from 3 independent replicates. The normalized average (thick lines) +/− SD (thin lines) of the carrier (red) and LMB-treated (yellow) cells are shown. Vertical dotted lines designate nuclear boundaries. **H**) Ca^2+^ chelation does not affect Chm7-GFP recruitment to the nuclear envelope in *apq12Δ* cells. Deconvolved inverted fluorescence micrographs of endogenously expressed Chm7-GFP in *apq12Δ* cells pretreated for 30 minutes with carrier (DMSO) or BAPTA-AM and shifted to the indicated temperature for 45 minutes. **I**) Plot quantifying the percentage of *apq12Δ* cells with Chm7-GFP foci at the indicated temperature and treatment. 3 independent replicates of > 100 cells were quantified per replicate. *P*-values are calculated from un-paired Student’s t-test where ns is *P* > 0.05.

Consistent with the possibility that Chm7 might be recognized by an export NTR, we identified two putative leucine-rich NESs at the C-terminus of Chm7 using the NES prediction algorithm, LocNES ((Xu et al., 2015); Figure 1A, B). Interestingly, the higher scoring predicted NES (NES2) overlapped with the potential MIM1 motif (Figure 1A, Figure 1 – figure supplement 1A). Indeed, the Chm7 putative MIM1 motif stands out from that of other budding yeast ESCRT-IIIs because of an additional isoleucine that contributes the fourth hydrophobic amino acid required for an effective type 1a NES (Figure 1 – figure supplement 1A). A second leucine in the middle of this region also aligns with the predictive spacing of residues in type 1b NESs (Figure 1-figure supplement 1A). Moreover, these putative NESs were conserved in Chm7 orthologues in other species of yeast, mice, flies and humans, although are curiously absent from *C. elegans* (Figure 1 – figure supplement 1B). As LocNES predicts NESs specific for the major export NTR, Xpo1/Crm1, we tested whether the inhibition of Xpo1 reversed the nuclear exclusion of Chm7-GFP. For these experiments, we took advantage of the *xpo1-T539C* allele, which sensitizes budding yeast Xpo1 to the Xpo1 inhibitor Leptomycin B (LMB)(Neville and Rosbash, 1999)(Figure 1E). As shown in Figure 1C, a 45 minute LMB treatment of cells expressing Chm7-GFP led to the striking accumulation of Chm7-GFP at the periphery of the nucleus, most often in highly fluorescent foci in over 60% of cells (Figure 1D), which is an underestimate as only midplanes were quantified.

The focal accumulation of Chm7-GFP at the nuclear periphery suggested that Chm7 was able to enter the nucleus upon Xpo1 inhibition. Therefore, to distinguish whether nuclear entry was driven by an active NTR-mediated nuclear import pathway, or, whether it was the result of passive diffusion across the NPC, we generated a Chm7-GFP fused to 5 maltose binding proteins (Chm7-MGM4); this ~280 kD protein would be extremely inefficient at transiting through the NPC unless it contained an NLS. Consistent with the idea that Chm7-GFP’s entry into the nucleus was governed by diffusion and not active NLS-mediated transport, the distribution of Chm7-MGM4 was indistinguishable from Chm7-GFP but, in contrast, incubation with LMB had no effect on its localization (Figure 1C-E).

As inhibition of Xpo1 led to Chm7-GFP accumulation at the nuclear envelope, we reasoned that it was likely that the prediction of NESs in Chm7 was likely accurate. However, as both NLS and NES prediction is of limited utility, we directly tested whether the predicted NESs were indeed sufficient to prevent an inert GFP reporter from accessing the nucleus. We first tested NES2 (Figure 1A). As shown in Figure 1F, a NES2_CHM7_-GFP fusion protein was only moderately excluded from the nucleus (middle panel) when compared to GFP alone (top panel, nucleus demarked using a dsRED-HDEL that localizes throughout the nuclear envelope-ER lumen) in a LMB-sensitive fashion, with average line profiles drawn from the cytosol and bisecting the nucleus suggesting an even distribution of fluorescence across the nuclear envelope (Figure 1G, red lines). Indeed, only upon testing a fragment containing both NES1 and NES2 (Figure 1A) could we observe a more obvious nuclear exclusion of this NES1-NES2_CHM7_-GFP construct that was sensitive to LMB treatment (Figure 1F, G; compare red and yellow lines). Therefore, it is likely that both predicted NESs contribute to the efficient export of Chm7. Consistent with this, the examination of a truncation of Chm7 (chm7_OPEN_; Figure 1A) lacking both NESs dramatically accumulates in one or two foci on the nuclear envelope in a way that is not impacted by LMB (Figure 1C, D and see (Webster et al., 2016)). Thus, the steady state nuclear exclusion of Chm7 in wildtype cells is determined by its passive diffusion into the nucleus and the Xpo1-mediated recognition of NESs in Chm7. That, having entered the nucleus, Chm7 accumulates in a focus along the nuclear periphery is consistent with its binding and activation at the INM. The latter being reflected in its focal accumulation, which would be consistent with a polymerization event.

That Chm7 can be recruited and activated at the INM without any perturbation to the nuclear envelope raises the possibility that there are no other upstream signals that are necessary to trigger Chm7 recruitment. While this is difficult to conclusively prove, we nonetheless tested whether Ca^2+^ could reflect an additional signal because of its role in other ESCRT-mediated membrane repair processes (Jimenez et al., 2014; Scheffer et al., 2014; Gong et al., 2017; Skowyra et al., 2018). As Chm7 is recruited to the nuclear envelope in *apq12Δ* cells when grown at elevated (37°C) temperatures (Webster et al., 2016), we evaluated whether this recruitment was influenced by chelating Ca^2+^ using BAPTA-AM. As shown in Figure 1H, there was no obvious change to the number of Chm7-GFP foci that appear during the temperature shift in the presence or absence of Ca^2+^(Figure 1H, I). Thus, it is unlikely that a Ca^2+^ signal is a major contributor to this pathway.

### Cytosolic exposure of the Heh1 WH domain is sufficient to recruit Chm7 to membranes

We hypothesized that Chm7 was excluded from the nucleus in order to prevent its untimely or inappropriate “activation” in the absence of a perturbation of the nuclear envelope barrier. Such a model predicts that there must be a nuclear binding partner that itself might be “hidden” from cytosolic Chm7; based on our and others’ prior work (Webster et al., 2014, 2016; Gu et al., 2017) the most obvious candidate was Heh1. To test this hypothesis, we generated deletion constructs of Heh1 coupled to the Red Fluorescent Protein (RFP) expressed behind the *GAL1* promoter (note that there is vacuolar autofluorescence even under repressed glucose conditions, see asterisks in Figure 2A). Unlike many other INM proteins that tend to back up into the ER upon overexpression (Lusk et al., 2007), Heh1-RFP continues to accumulate at the INM even at high levels due to its use of an active NTR-dependent INM targeting pathway (See Figure 2A, B and (King et al., 2006)). Thus, even when overexpressed at levels that we estimate to be an order of magnitude higher than endogenous levels, the majority of Heh1 is localized to the INM and would be predicted to be inaccessible to cytosolic Chm7 (Figure 2A, galactose, middle panels). Consistent with this, we observed no change to the steady state distribution of endogenously-expressed Chm7-GFP, which includes a minor fraction within a nuclear envelope focus in ~30% of cells (Webster et al., 2016; Figure 2A). Deletion of the LEM domain of Heh1 also had no effect on Chm7-GFP distribution as heh1(51-834)-RFP was also exclusively localized at the INM (Figure 2A).

**Figure 2:**
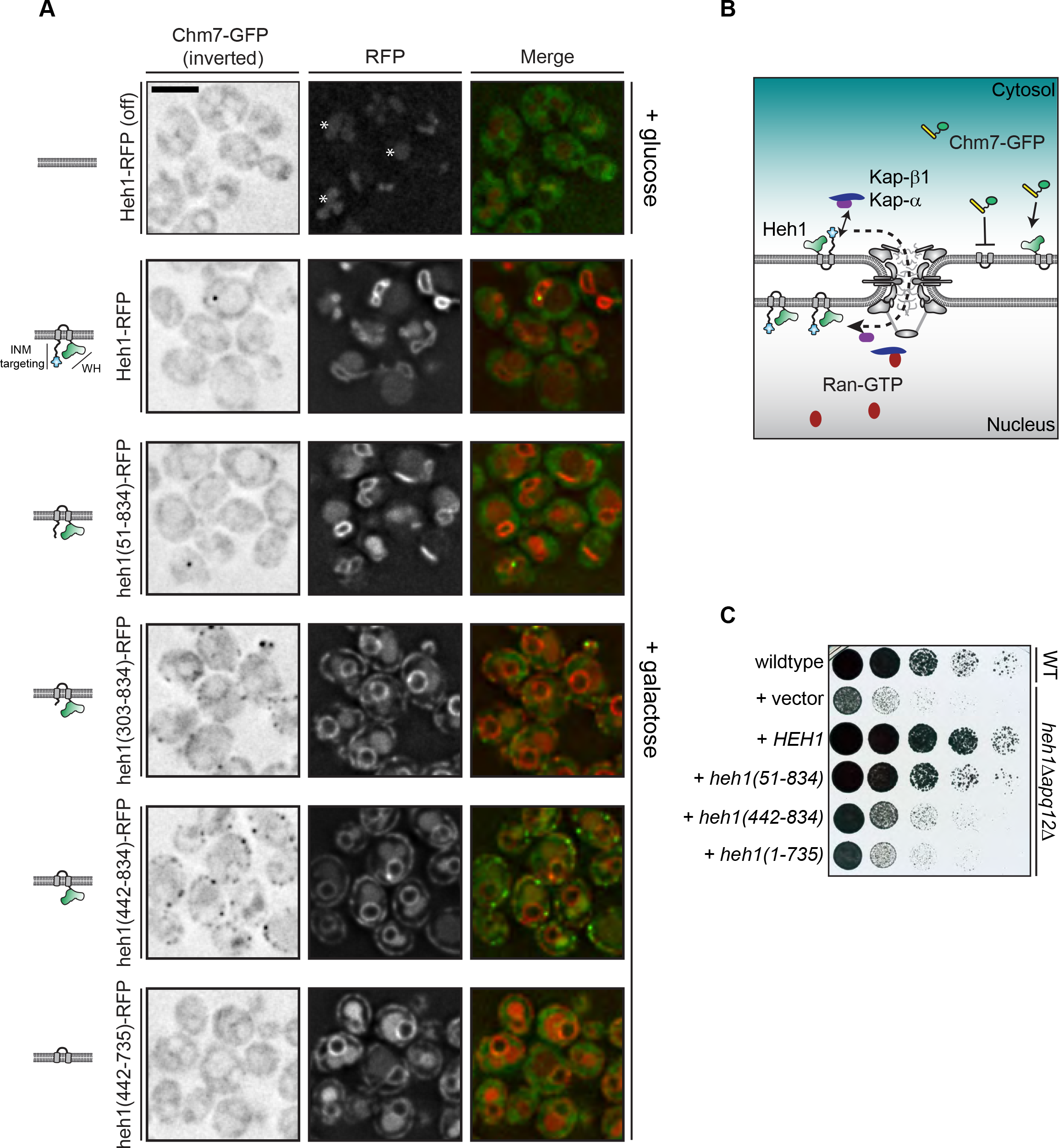
Cytosolic exposure of the Heh1 WH domain is sufficient to recruit Chm7 to ER membranes. **A**) Deconvolved fluorescence micrographs of endogenously expressed Chm7-GFP (inverted) either prior to (+glucose) or after 2 hours of overexpression (+galactose) of RFP-tagged full length and truncations of Heh1 (depicted in cartoons at left in a lipid bilayer). Asterisks indicate vacuolar autofluorescence in the red channel. Scale bar is 5 μm. **B**) Cartoon of experiment and interpretation of A. The efficient INM targeting of Heh1 depends on Kap-α/Kap-β1 (blue and purple) and on Ran-GTP (red). **C**) Tenfold serial dilutions of the indicated strains spotted onto YPG plates to express the indicated truncations of Heh1. Plates imaged after growth at RT for 48 h.

We next tested deletions that encompassed the putative NLSs in Heh1 including heh1(303-834), and heh1(442-834), which resulted in the accumulation of these truncations throughout the cortical ER. Strikingly, we observed a concurrent re-distribution of Chm7-GFP into foci that colocalized with the RFP signal (Figure 2A). In the case of heh1(442-834), only the WH domain is available for Chm7 binding. Consistent with this, there was a complete lack of Chm7-GFP at the ER in cells expressing heh1(442-735), where the WH is removed (Figure 2A). Thus, exposure of the Heh1 WH domain to the cytosol is both necessary and sufficient to recruit Chm7-GFP to ER membranes.

We next assessed the functional importance of the Heh1-WH domain to *apq12Δ* cells, which require both *CHM7* and *HEH1* for full viability (Yewdell et al., 2011; Bauer et al., 2015; Webster et al., 2016)(Figure 2C). Interestingly, the loss of fitness observed in *heh1Δapq12Δ* cells could only be rescued by the gene encoding full length Heh1 or the *heh1(51-834)* allele. In contrast, deletions that resulted in Heh1 mistargeting or those that are unable to recruit Chm7 (e.g. *heh1(1-735),* which lacks the coding sequence for the WH domain) were unable to fully complement growth. Thus, while the WH domain is important, the N-terminal INM targeting domain is also a critical component of Heh1 functionality in the context of *apq12Δ* cells.

### Chm7 binds to an INM platform

The localization data clearly pointed to a direct interaction between the WH domain of Heh1 and Chm7. Unfortunately, we were unable to detect a stable interaction *in vitro* with purified recombinant proteins (one example shown in Figure 3 – figure supplement 1A), although we note such an interaction has been shown with the human versions of these proteins (Gu et al., 2017). While there are many potential reasons for these negative data, one possibility is that there are additional proteins (or lipids) that contribute to the interaction *in vivo*. To test this idea, we affinity purified endogenously expressed Chm7-GFP from whole cell extracts using anti-GFP nanobody-coupled beads (Figure 3 – figure supplement 1B) and subjected protein eluates to MS/MS peptide identification. Consistent with the idea that Chm7 is localized throughout the cytosol in a potentially inactive form, we detected few specific peptide spectra with the exception of Chm7 itself when compared to proteins derived from wildtype cell extracts that bind non-specifically to the anti-GFP beads (Figure 3A). To facilitate visualization, we directly relate the average spectral counts (two experiments) from bound fractions of affinity purifications of Chm7-GFP and no-GFP controls in Figure 3A.

**Figure 3:**
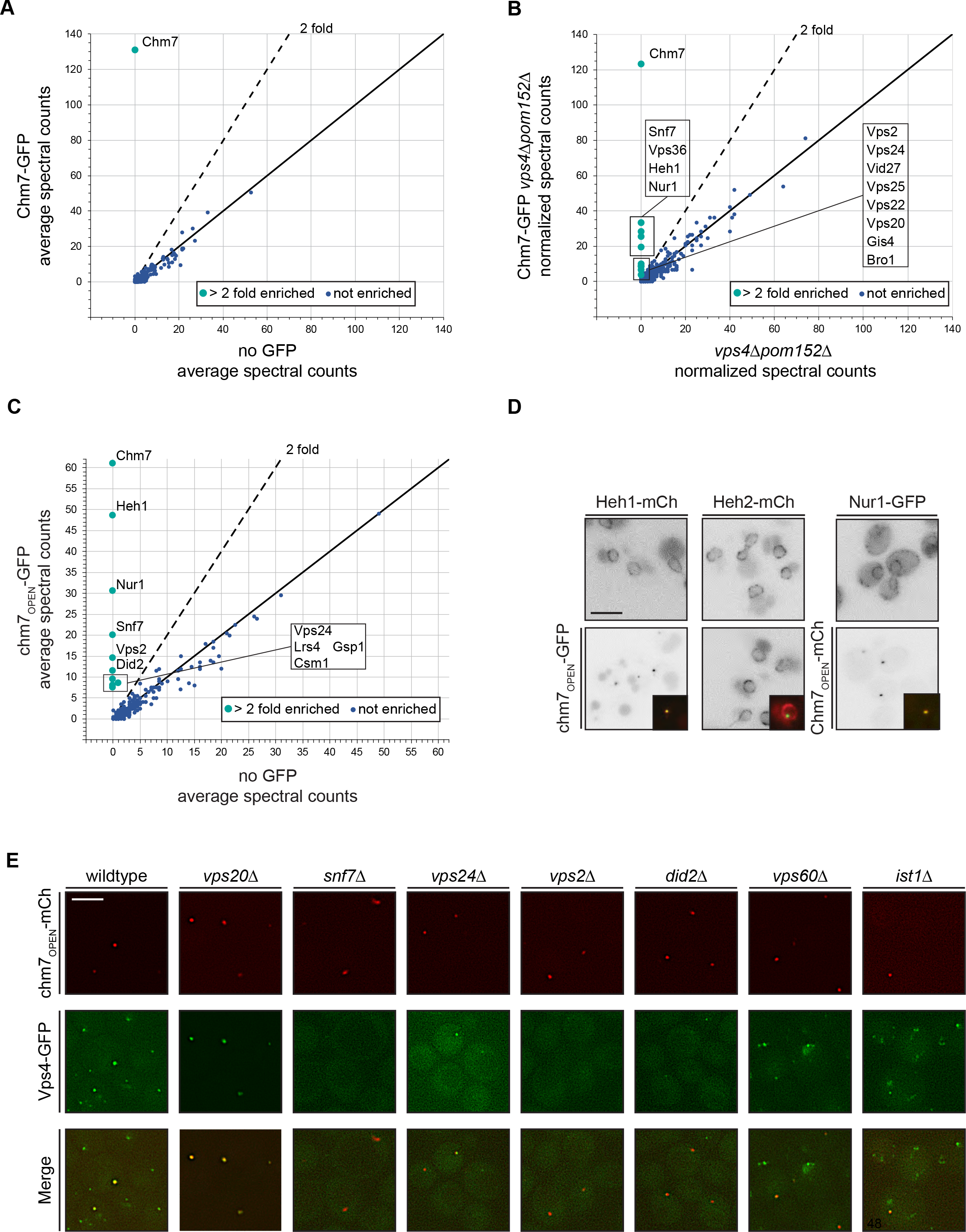
Chm7 binds to Heh1, Nur1 and downstream ESCRTs required for Vps4 recruitment. **A-C**) Affinity purifications of endogenously expressed Chm7-GFP and chm7_OPEN_-GFP were performed from wildtype and *vps4Δpom152Δ* cells. Bound proteins were eluted and subjected to LC-MS/MS peptide identification. Scatter plots of the number of peptide spectra identifying the indicated proteins were directly compared to those identified in “no GFP” samples. Dotted line and teal circles represent peptides found at least 2 fold enriched over non-specific proteins (blue). Plots A and C represent an average of two replicates of normalized spectral counts of peptides identified by MS. **D**) Heh1 and Nur1 but not Heh2 colocalize with chm7_OPEN_. Deconvolved inverted fluorescence micrographs of Heh1-mCherry, Heh2-mCherry, with and without chm7_OPEN_-GFP, and Nur1-GFP with and without chm7_OPEN_-mCherry. Merge of green and red channels shown in inset. **E**) Vps4 recruitment to chm7_OPEN_ requires Snf7 and downstream ESCRTs. Deconvolved fluorescence micrographs of Vps4-GFP and chm7_OPEN_ -mCherry in the indicated strain backgrounds. Green, red and merged images shown.

We therefore turned to examining the interactome of Chm7-GFP under conditions in which it accumulates at the nuclear envelope, for example in *vps4Δpom152Δ* cells, which we had previously shown leads to Chm7-GFP accumulation within a nuclear envelope domain enriched for malformed NPCs (Webster et al., 2014, 2016). Shot-gun MS identification of peptides derived from bound proteins to Chm7-GFP now revealed specific interactions with several proteins including the ESCRT-III, Snf7, the ESCRT-II, Vps36 (the ESCRT-III Vps20 was also found, but with comparably low abundance using this semi-quantitative approach; Figure 3B). Most interestingly, dozens of spectra specific for Heh1 were identified. Considering Heh1 is a low abundant integral membrane protein (measured to be as low as 428 molecules/cell; (Kulak et al., 2014)), this result was particularly striking. In addition, another low abundant (~354 molecules/cell; (Kulak et al., 2014)) integral INM protein, Nur1 was also detected. As Nur1 is known to interact with Heh1 within the CLIP (chromosome linkage INM proteins) complex (Mekhail et al., 2008), these data suggest that Chm7 engages Heh1 within a broader INM platform, at least in the context of cells lacking *VPS4*. Of note, no components of the NPC were specifically detected, nor was Heh2.

We next tested binding partners of chm7_OPEN_-GFP (Figure 3 – figure supplement 1B), which also provides a potential mimic of the physiological circumstances when Chm7 is recruited to the nuclear envelope. In this case, Heh1 was the top hit (Figure 3C). In addition to Nur1, other members of CLIP were also specifically identified including Lrs4 and Csm1. Curiously, Gsp1 (budding yeast Ran) was also found (Figure 3C). Further, alongside Snf7, other ESCRT-IIIs including Vps2, Vps24 and Did2 were detected (Figure 3C). In contrast to bound proteins purified with Chm7-GFP in the *vps4Δpom152Δ* cells, we did not detect any specific peptides for ESCRT-II subunits or Vps20. We surmise this is likely because Chm7-GFP can also be seen in cytosolic foci in *vps4Δ* cells (See (Webster et al., 2016) and Figure 5A), whereas chm7_OPEN_-GFP exclusively localizes to the nuclear envelope. As a further test of the specificity of the interactions between chm7_OPEN_-GFP and integral INM proteins, we observed the near-quantitative accumulation of both Heh1 and Nur1 fluorescent fusion proteins (produced at endogenous levels) at the chm7_OPEN_ focus, while the distribution of Heh2 was unaltered (Figure 3D).

**Figure 4:**
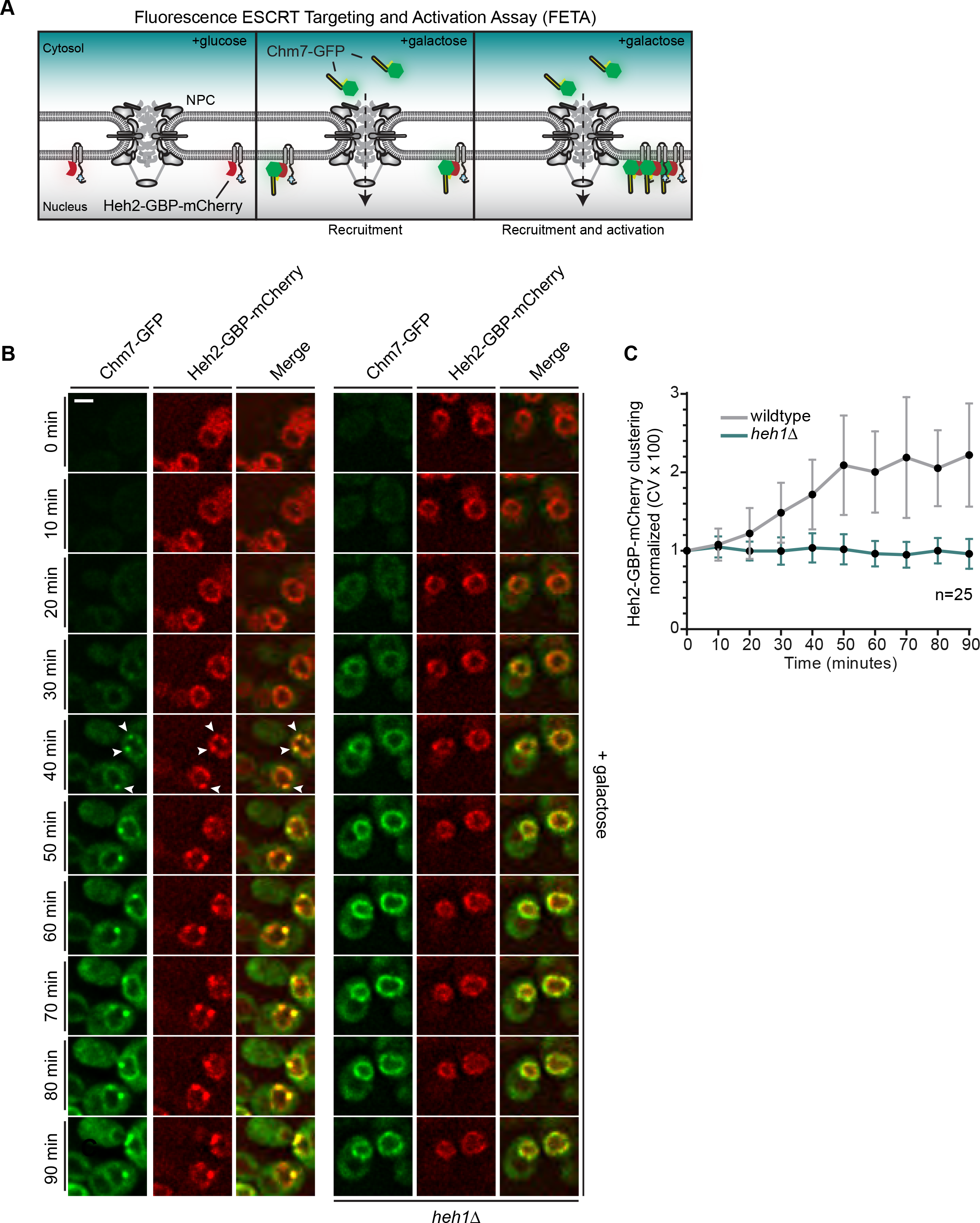
Heh1 is required to induce the focal accumulation of Chm7. **A**) Schematic of “Fluorescence ESCRT Targeting and Activation Assay” where Heh2 is expressed as a fusion to GFP binding protein (GBP; red). Chm7-GFP is expressed by the addition of galactose to the growth medium. The focal accumulation of Heh2-GBP-mCherry is interpreted as Chm7 activation. **B**) Deconvolved fluorescence micrographs of overexpressed Chm7-GFP and Heh2-GBP-mCherry at the indicated timepoints after addition of galactose to the growth medium. Scale bar is 5 μm. **C**) As a metric for the clustering of Heh2-GBP-mCherry, a coefficient of variation (CV) of the Heh2-GBP-mCherry fluorescence along the nuclear envelope in a mid-section was calculated over time. Mean and SD normalized to 0 timepoint are shown. n=25.

**Figure 5:**
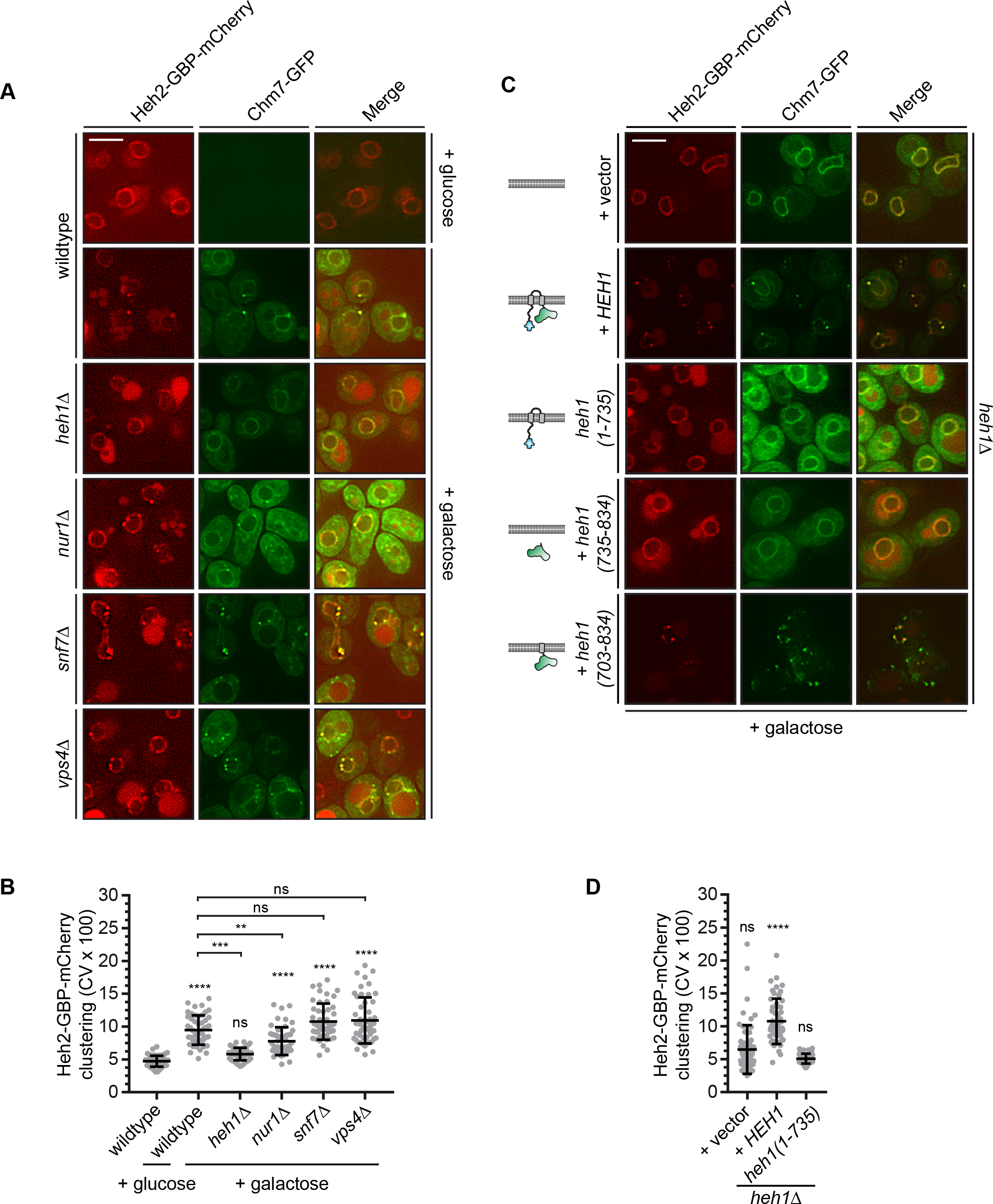
The Heh1 WH domain and a transmembrane anchor are necessary and sufficient for Chm7 activation. **A**) Deconvolved fluorescence micrographs of Heh2-GBP-mCherry and overexpressed Chm7-GFP in the indicated genetic backgrounds, either prior to (glucose) or after 90 minutes in galactose to drive Chm7-GFP production. Green and red channels with merge shown. Scale bar is 5 μm. **B**) Plot of the CV of Heh2-GBP-mCherry fluorescence (as in Figure 4C) in the indicated strains before (glucose) or after 90 minutes of galactose addition to induce Chm7-GFP production. Data is from three independent replicates where 50 cells/genotype/replicate were counted. *P*-values are from two-way ANOVA with Dunnett’s test where ns is *P* > 0.05, ** *P* ≤ 0.01, *** *P* ≤ 0.001, and **** *P* ≤ 0.0001. **C**) Deconvolved fluorescence micrographs of Heh2-GBP-mCherry and overexpressed Chm7-GFP produced for 90 minutes in galactose. All images are from *heh1Δ* strains expressing the indicated genes encoding *HEH1* and several deletion constructs (schematized at left in a lipid bilayer). **D**) Plot of the CV of Heh2-GBP-mCherry fluorescence (as in Figure 4C) in an *heh1Δ* strain expressing *HEH1* or *heh1(1-735)* after 90 minutes of galactose addition to induce Chm7-GFP production. Data is from three independent replicates of 50 cells per strain. *P*-values are from two-way ANOVA with Dunnett’s test where ns is *P* > 0.05, **** *P* ≤ 0.0001.

We noted that although we detected several additional ESCRT-III proteins in the affinity purifications of chm7_OPEN_-GFP, we did not detect any peptides for Vps4. Thus, we investigated whether a functional Vps4-GFP fusion (Adell et al., 2017) could also be specifically recruited to the chm7_OPEN_ focus at the nuclear envelope, which it was in virtually all cells (Figure 3E). The fluorescence intensity of Vps4-GFP could be correlated to that of the chm7_OPEN_-mCherry focus, suggesting a close relationship between the number of Chm7 and Vps4 molecules recruited to this nuclear envelope site (Figure 3 – figure supplement 1C). This result provided the opportunity to explore the determinants of Vps4 recruitment at the nuclear envelope by deleting the genes encoding ESCRT subunits found in the chm7_OPEN_-GFP affinity purifications including *SNF7*, *VPS24, VPS2*, and *DID2*. In all cases, Vps4-GFP recruitment to Chm7_OPEN_-mCherry foci was reduced or, in the case of *snf7Δ* completely eliminated, with minimal impact on the accumulation of chm7_OPEN_-mCherry itself (Figure 3E; Figure 3 – figure supplement 1D, E); although we noted that the average area encompassed by the chm7_OPEN_-mcherry focus was increased in *snf7Δ* cells (Figure 3E; Figure 3 – figure supplement 1F). Of further interest, the deletion of *VPS20* not only abolished Vps4-GFP localization in cytosolic puncta, but in fact led to a ~3 fold increase of Vps4-GFP at the nuclear chm7_OPEN_ focus (Figure 3E, Figure 3 – figure supplement 1D) while the total Vps4-GFP protein levels were not notably altered in any of the ESCRT deletion backgrounds (Figure 3 – figure supplement 1G). Together, these data support a model in which Vps4 can be recruited to the INM, likely through interactions with ESCRT-III subunits recruited alongside or downstream of Snf7.

### Fluorescence ESCRT Targeting and Activation (FETA) Assay

Interestingly, as shown in Figure 3D, chm7_OPEN_ is able to shift the distribution of both Heh1 and Nur1 from an evenly-distributed nuclear peripheral localization to one that is co-localized with the chm7_OPEN_ focus. This raised the formal possibility that Chm7 recruitment to the nuclear envelope might in fact be independent of and precede binding to Heh1; in such a model Heh1 would be required for its focal accumulation, which we interpret to be “activation”. Thus, this result illustrated the need to develop a better controlled experimental system where the mechanism Chm7 recruitment and activation can be decoupled. We thus generated an experimental approach that we termed the <u>F</u>luorescent <u>E</u>SCRT <u>T</u>argeting and <u>A</u>ctivation Assay (FETA; Figure 4A). FETA exerts both temporal control over the expression of Chm7-GFP (through the *GAL1* promoter) in addition to spatial control over its recruitment to the nuclear envelope by binding to a GFP-nanobody (GFP-binding protein/GBP) appended to Heh2. Thus, without any perturbations to the nuclear envelope barrier, we can monitor the recruitment and activation of Chm7-GFP at the nuclear envelope, the latter of which we interpret as the local clustering of Chm7 as the most logical visual outcome of Chm7 polymerization at the level of fluorescence microscopy.

By shifting cells to medium containing galactose, we induced the expression of Chm7-GFP and monitored its distribution by timelapse microscopy. As shown in Figure 4B, Chm7-GFP was first observed in the cytosol but accumulated at the nuclear envelope within 20 minutes. Importantly, this nuclear envelope binding was due to its interaction with Heh2-GBP-mCherry as overexpression of Chm7-GFP in strains lacking Heh2-GBP-mCherry (Figure 1C) or lacking GBP (Figure 4 – figure supplement 1A, B) did not lead to nuclear envelope accumulation or clustering. In contrast, Heh1-mCherry was incorporated into the Chm7-GFP-Heh2-GBP foci (Figure 4 – figure supplement 1C). Interestingly, nearly simultaneously with the broader nuclear envelope-localization, Chm7-GFP and Heh2-GBP-mCherry accumulated in multiple foci throughout the nuclear envelope (see arrowheads at 40 min). These foci coalesced into one or two foci/cell over the length of the timecourse (90 min; Video 1). As a means to quantify this focal accumulation, we calculated a coefficient of variation (CV) of the mCherry fluorescence along the nuclear envelope in a mid-plane, which we plotted over time (Figure 4C). This approach faithfully represented the observed clustering, which reached a maximum value between 50 and 60 minutes (Figure 4B, C).

With the ability to temporally resolve recruitment and “activation,” we next interrogated how Heh1 impacted these steps. Strikingly, the induction of Chm7-GFP expression in *heh1Δ* cells lead to its accumulation at the nuclear envelope at a similar timepoint as in wildtype cells, however, we did not observe any focal accumulation with a CV remaining at ~1 over the 90 minute timecourse (Figure 4B, C, and Video 1). Thus, Chm7 recruitment to the INM is not sufficient to lead to Chm7-GFP clustering. Instead, activation requires Heh1.

We next investigated whether other Chm7-interacting partners influenced Chm7-GFP clustering in the FETA assay beginning with Nur1. Interestingly, deletion of *NUR1* led to a statistically-significant drop in the CV of Heh2-GBP-mCherry at the end point of the FETA assay suggesting it could also contribute to Chm7 activation (Figure 5A, B). Consistent with this idea, we also observed considerably less chm7_OPEN_-GFP accumulation at the nuclear envelope in *nur1Δ* cells (Figure 5 – figure supplement 1A, B). However, we also noted that the total levels of Heh1 are reduced in *nur1Δ* cells (Figure 5 – figure supplement 1C), suggesting that this effect may be indirect and ultimately mediated through Heh1. We further investigated the impact of deleting both *SNF7* and *VPS4* on the extent of Heh2-GBP-mCherry clustering in the FETA assay. In both cases, the CV of Heh2-GBP-mCherry was unaltered (Figure 5B). Yet, qualitatively, we observed that there were more discrete foci at the nuclear envelope in both of these genetic backgrounds, in addition to some foci (lacking Heh2-GBP-mCherry) in the cytoplasm (Figure 5A). Thus, these downstream components could impact other factors that elude measurement and interpretation with this approach.

The lack of Heh2-GBP-mCherry clustering in *heh1Δ* cells also provided a genetic background to more fully vet the domains of Heh1 that contribute to Chm7 activation. First, introduction of *HEH1* on a plasmid rescued clustering of Chm7-GFP and Heh2-GBP-mCherry in the *heh1Δ* strains, confirming that lack of clustering was indeed due to the absence of Heh1 (Figure 5C, D). In contrast, expression of *heh1(1-735)*, which lacks the C-terminal WH domain, failed to restore Chm7 focal accumulation and Heh2-GBP-mCherry clustering suggesting that the WH domain was required for Chm7 activation (Figure 5C, D). Interestingly, however, the WH domain alone was insufficient to rescue clustering but instead it required a membrane anchor through a Heh1-transmembrane domain (Figure 5C, bottom panel). In the latter case, we also observed Chm7-GFP foci throughout the cytosol as the heh1(703-834) construct does not contain INM targeting sequences, consistent with data presented in Figure 2A. Thus, there is a clear coupling between the Heh1 WH domain and the membrane required for Chm7 activation.

### Chm7-Heh1 interactions drive nuclear envelope herniation and expansion

Our data support a model in which Chm7 and Heh1 are spatially segregated, but, upon binding Chm7 is locally activated at a membrane interface. To investigate how Chm7 activation could translate into a mechanism capable of sealing a nuclear envelope hole, we turned to a correlative light EM (CLEM) approach to investigate nuclear envelope morphology at sites of Chm7 activation. While our initial attempts focused on examining sites of Chm7-GFP localization in NPC assembly-defective strains that have nuclear envelope herniations like in *apq12Δ* and *nup116Δ* cells, the combination of the low abundance (and transience) of Chm7-GFP at these nuclear envelope foci coupled to the loss of GFP fluorescence through the freeze-substitution process precluded this as a viable approach. Thus, again, we turned to chm7_OPEN_-GFP (and Chm7-GFP in *vps4Δpom152Δ* cells) as proxies to interrogate the membrane morphology at sites of surveillance (hyper) activation.

As shown in in panel *i* of Figure 6A and B, we could effectively correlate fluorescence images and electron tomograms (Kukulski et al., 2012), which showed the chm7_OPEN_-GFP foci apposed to the INM. Remarkably, this fluorescence demarked extensions of the INM that invade the nucleus and form a fenestrated network of INM cisterna – the lumen of the cisterna is continuous with the lumen of the nuclear envelope and is colored teal or purple to facilitate visualization. In Figure 6A and B, the panels *ii-iv* represent slices along the Z axis of the tomograms; 3D models were generated by isosurface rendering, which can be visualized as still frames (Figure 6A, B, *v-viii*) and in movies (Videos 2 and 3). These 3D views facilitate the observation of nuclear pores (denoted by stars) in addition to a perspective of the extent of the network of fenestrated cisternal membranes at the INM. Nearby, fenestrated ER membranes also appeared with intriguing frequency (Figure 6A, *vii, viii*; 6B, *iii*; 6 out of 14 tomograms).

**Figure 6:**
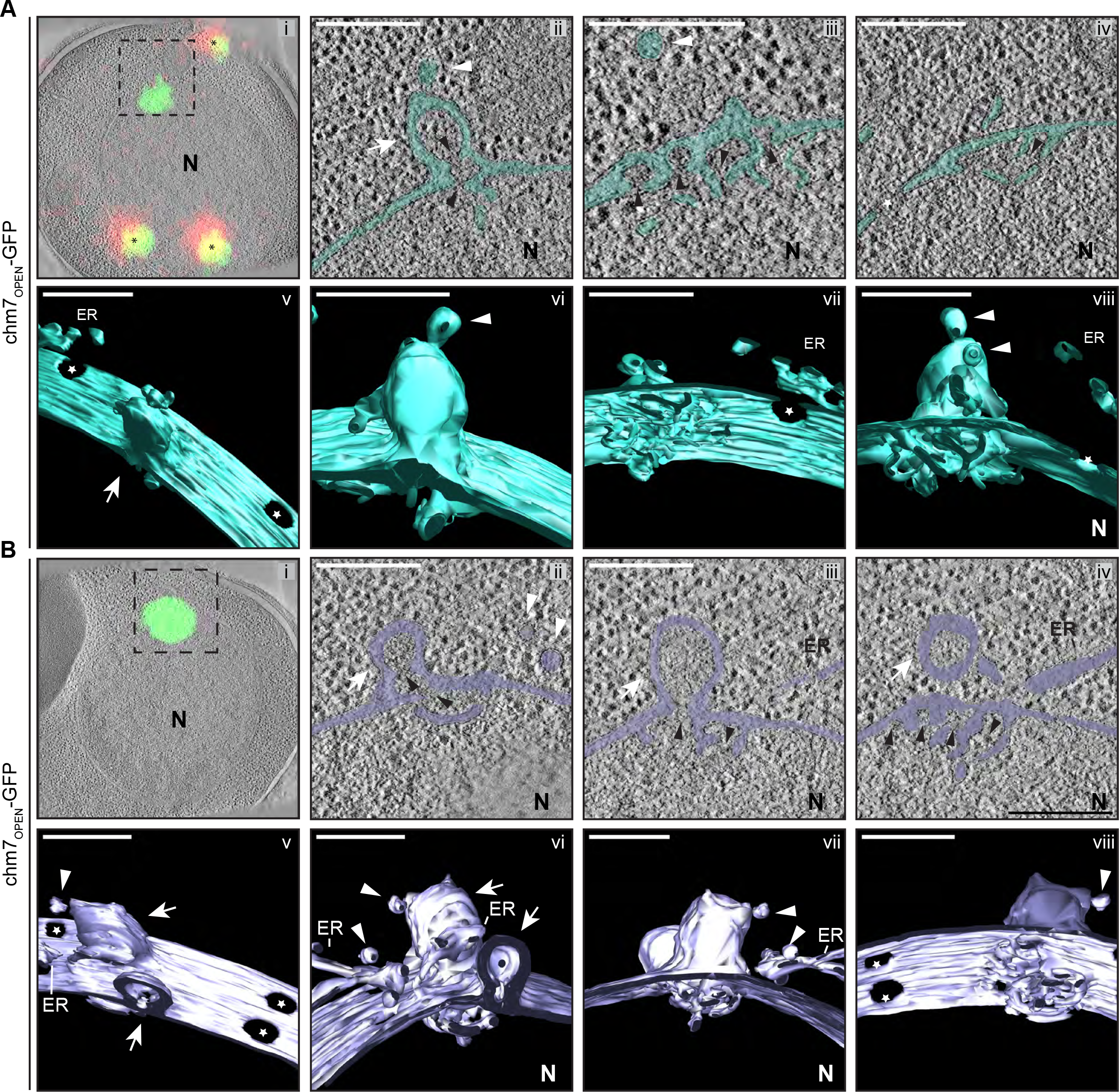
chm7_OPEN_ associates with a network of intranuclear fenestrated cisterna often below nuclear envelope herniations. **A**, **B**) Correlative light and electron microscopy of 300 nm thick sections was used to examine the morphology of the nuclear envelope at sites of chm7_OPEN_-GFP accumulation. *i.* Overlay of fluorescent and electron micrographs with tetrafluorescent fiducials used for correlative alignment marked with (*); boxed region is magnified in *ii-iv*. *ii-iv.* Several views along the z-axis of the tomogram shown with the nuclear envelope/ER lumen filled with teal or light purple. *v-viii*. 3D models were generated and several perspective views are shown. White arrows are herniations, while arrowheads vesicles, black arrowheads are constrictions or necks of budding herniations, stars are nuclear pores. N is nucleus. Scale bars are 250 nm.

The INM-associated cisternal network was often (11 out of 14 tomograms) found underneath balloon-like herniations of the nuclear envelope (i.e. both the INM and ONM) that extended several hundred nanometers into the cytosol (Figure 6A, B). The lumen of the herniations were open to the nucleoplasm and tapered into ~45 nm-diameter membrane “necks”: in the image shown in Figure 6A, *ii*, two “necks” can be observed (black arrowheads). Often (in 10 out of 14 tomograms) vesicles appear nearby the sites of herniations some of which can be seen either fusing with, or fissioning from, the ONM (Figure 6, white arrowheads). Additional INM evaginations (extending into the nuclear envelope lumen, black arrowheads) are observed in tandem arrays that are suggestive of precursors of the herniations (Figure 6A, *iii*). Indeed, often several nuclear envelope herniations can be observed in a single tomogram, as indicated by white arrows in Figure 6B. Interestingly, these membrane deformations could also be formed in the absence of *SNF7* (Figure 6 – figure supplement 1A, B, Video 4).

Similar membrane morphology was also observed when CLEM was applied to the focal accumulation of Chm7-GFP in *vps4Δpom152Δ* cells (Figure 7A, *i,*). Consistent with the idea that the INM expansion and nuclear envelope herniation need not occur simultaneously, in Figure 7A an example of a single herniation (with two necks; see black arrow heads) is shown. A nuclear-perspective (bottom-up, Figure 7, *v*) view allows a direct comparison between the herniation neck at the Chm7-GFP signal and nuclear pores that would be filled with NPCs (stars). Lastly, a more dramatic example of a *vps4Δpom152Δ* nuclear envelope where connections between the INM and a cisternal, lamellar membrane with multiple deformations is presented in the Figure 7B. Interestingly, in this thick section, no nuclear envelope herniation is observed suggesting that there is no implicit link between the INM network and nuclear envelope herniation; these two morphologies might arise stochastically and are not necessarily directed in one, or the other, direction. Taken together, these data support a model in which the interaction of Chm7 and Heh1, and its activation, can lead to expansion of the INM and the formation of nuclear envelope herniations.

**Figure 7:**
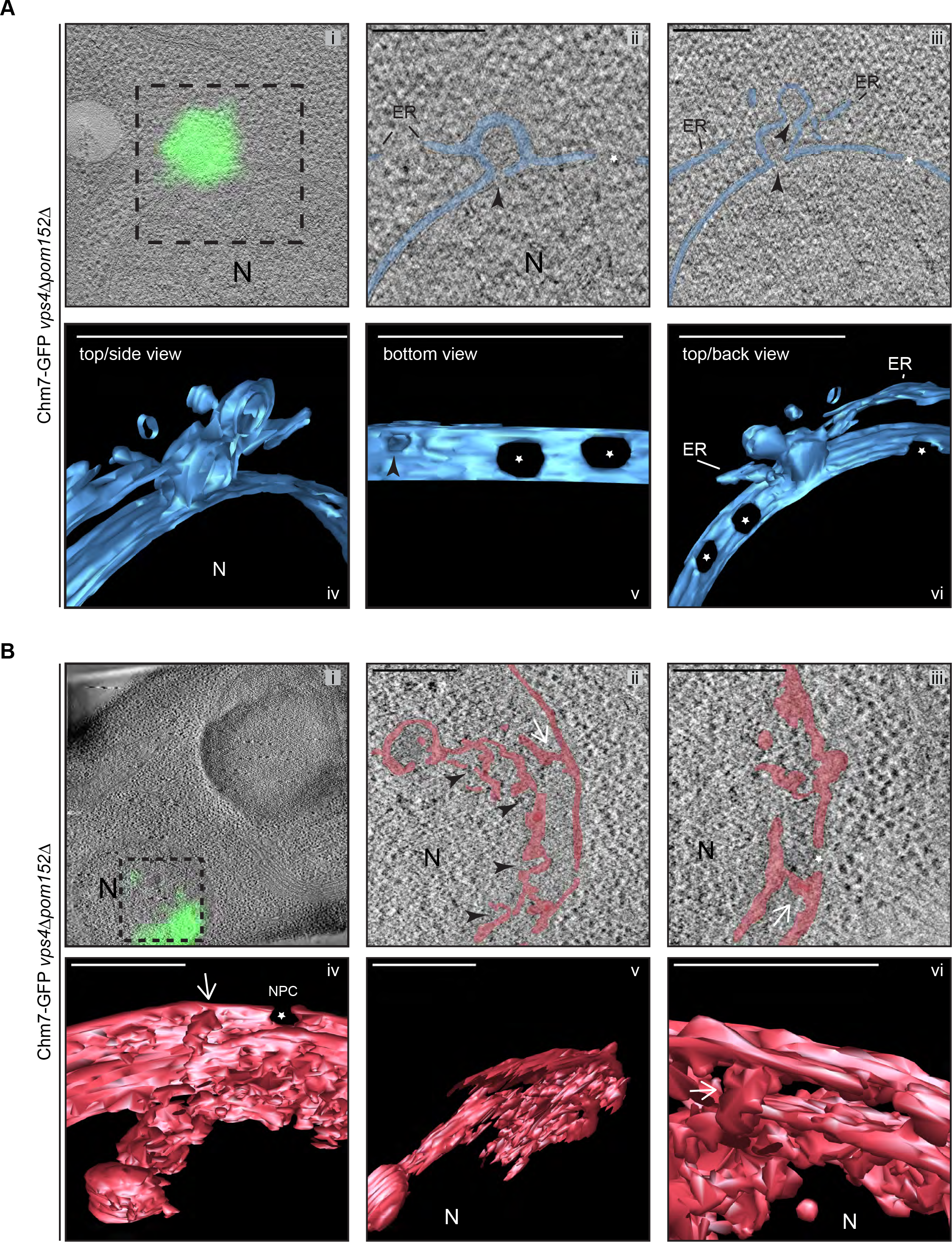
Chm7 associates with nuclear envelope herniations and intranuclear membranes. **A, B**) Correlative light and electron microscopy of 300 nm thick sections was used to examine the morphology of the nuclear envelope at sites of Chm7-GFP accumulation in *vps4Δpom152Δ* cells. *i.* Overlay of fluorescent and electron micrographs; boxed region is magnified in *ii* and *iii*. *ii, iii.* Several views along the z-axis of the tomogram shown*. iv-vi*. 3D models were generated with indicated perspective views shown. Nuclear envelope/ER lumen colored blue or red. Black arrowheads are herniation necks or sites of membrane constriction, stars are nuclear pores. N is nucleus. Scale bars are 250 nm.

### Morphologically distinct nuclear envelope herniations are associated with defects in NPC biogenesis

It was tempting to speculate that the nuclear envelope herniations that we observed under conditions of Chm7 activation were directly analogous to those observed in genetic backgrounds where NPC assembly is perturbed, like in *apq12Δ* and *nup116Δ* backgrounds. To perform a direct comparison, we first confirmed that, as in *nup116Δ* cells (Wente and Blobel, 1993), NPC-like structures were found at the bases of the nuclear envelope herniations seen in cells lacking *APQ12* (Scarcelli et al., 2007) by staining thin sections with the MAb414 antibody that recognizes several FG-nups. As shown in Figure 8A, gold particles that label the MAb414 antibody were specifically found at the bases of these herniations confirming that they emanate from structures with nups. Furthermore, the diameter of the bases of these herniations averaged 78 nm, which while statistically similar to those found at mature NPCs (mean of 87 nm), were considerably larger than the ~45 nm diameter openings found at the necks of herniations caused by Chm7 (Figure 8B, Video 5). Lastly, the lumen of the herniations associated with both *apq12Δ* (Figure 8A, C) and *nup116Δ* cells (Figure 8 – figure supplement 1A, Video 6 and (Wente and Blobel, 1993)) are filled with electron density, whereas those associated with Chm7 appear to be empty (Figure 6A, B). Thus, we suggest that the herniations associated with overactive Chm7 and those associated with NPC assembly are morphologically distinct.

**Figure 8:**
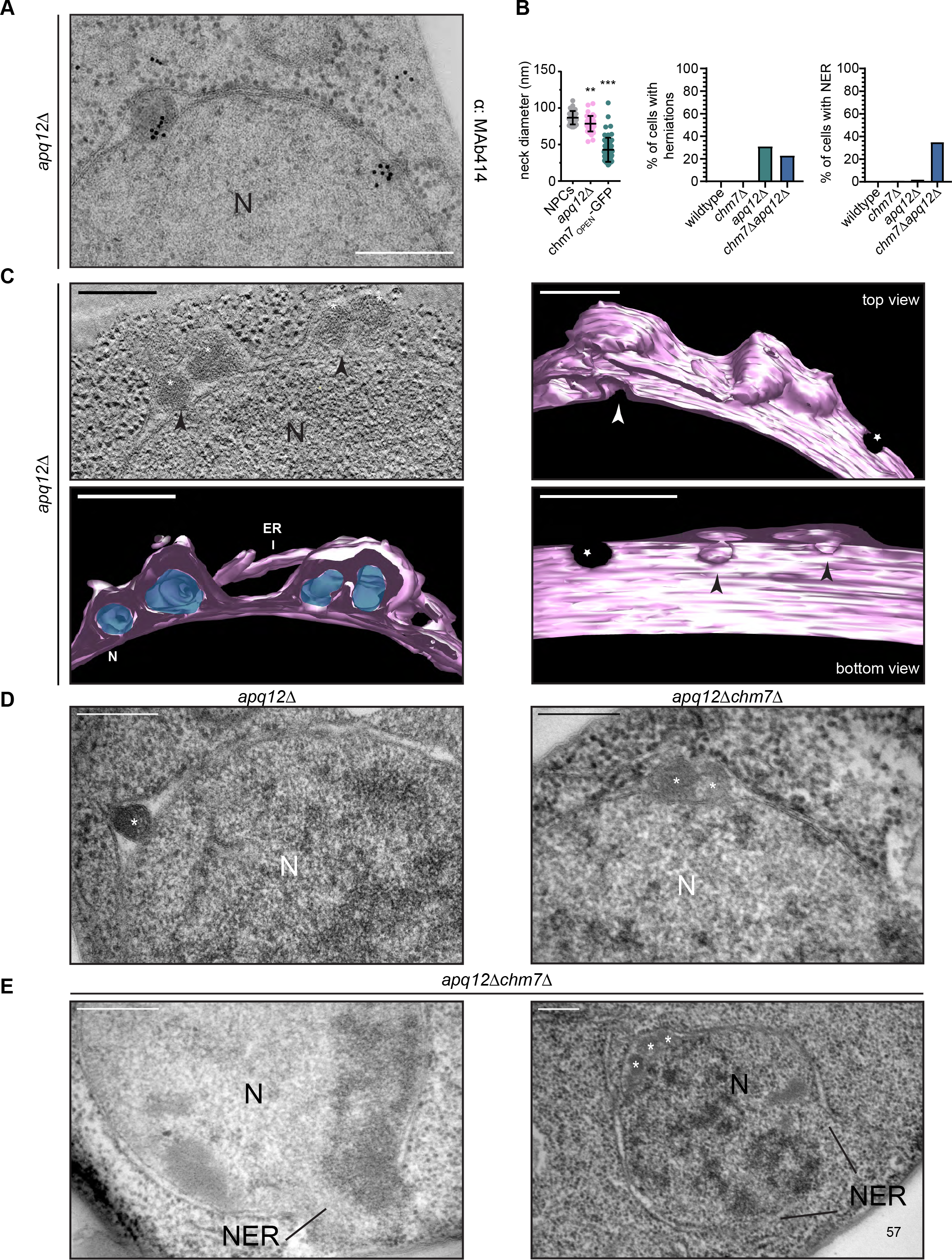
*CHM7* is required to maintain the integrity of the nuclear membranes in the context of nucleoporin-associated herniations. **A**) Nucleoporins are found at the bases of nuclear envelope herniations in *apq12Δ* cells. Immunogold labelling of thin sections of *apq12Δ* cells with 5 nm gold-conjugated secondary antibodies that detect MAb414 labeled nups at bases of herniations. **B**) Left: Plot of the diameter of herniation necks in the indicated genetic backgrounds and fenestrations within the intranuclear membrane network associated with chm7_OPEN_. Middle: Plot of the percentage of nuclei of the indicated strains where nuclear envelope herniations are observed. Right: Plot of the percentage of nuclei in the indicated strains where nuclear envelope ruptures (NER) are observed. At least 100 cells from each genotype were quantified. *P*-values are from Student’s t-test, where ** *P* ≤ 0.01, *** *P* ≤ 0.001. **C**) Electron tomograph of 300 nm thick section of *apq12Δ* cells grown at 37°C for 2 hours prior to high pressure freezing. Note the electron density within the herniations. Tomograms were segmented to generate a 3D model with perspective views shown (membranes/nuclear envelope-ER lumen colored pink with electron density within herniation blue). Arrowheads point to herniation necks and stars are nuclear pores. Scale bars are 250 nm. **D**) Nuclear envelope herniations associated with nups persist in the absence of *CHM7*. Representative electron micrographs of the *apq12Δ* and *chm7Δapq12Δ* strains grown for 2 hours at 37°C prior to high pressure freezing. Asterisks denote herniation lumen. N is nucleus. **E**) Nuclear envelope ruptures (NER) are observed in *chm7Δapq12Δ* cells. Electron micrographs of *chm7Δapq12Δ* depicting nuclear envelope ruptures (NER). Nucleus is indicated with “N”. Scale bars are 250 nm.

That activated Chm7 might drive membrane expansion and nuclear envelope herniations that have unique characteristics to those found in NPC assembly mutants does not exclude the possibility that Chm7 might nonetheless contribute to the formation of both of these herniation types. We therefore next investigated whether deletion of *CHM7* impacted the prevalence of herniations in *apq12Δ* cells. As shown in Figure 8B and D, deletion of *CHM7* had little impact on the number of herniations observed in thin sections of *apq12Δ* cells, which were only modestly reduced (23% versus 31% of nuclei; Figure 8B). Most strikingly, however, we observed that 35% of the nuclei in the thin sections of *apq12Δchm7Δ* cells (Figure 8B, E and Figure 8 – figure supplement 2) had large (>500 nm) discontinuities in their nuclear membranes, which suggests that these nuclei were unstable and could rupture. Similar nuclear envelope discontinuities were observed in *apq12Δsnf7Δ* strains (Figure 8 – figure supplement 3). Indeed, in some cases we could observe nucleoplasm escaping into the cytosol (Figure 8E, left panel). This result provides an explanation for the striking loss of NLS-GFP reporter accumulation in the nucleus that was observed in only ~35% of *apq12Δchm7Δ* cells (Webster et al., 2016). Thus, while these NPC-assembly-associated nuclear envelope herniations might not require *CHM7* or *SNF7* for their biogenesis, ESCRTs are nonetheless required to maintain the integrity of the nuclear membranes in the context of these herniations.

## Discussion

While there has been considerable focus over the last few decades on mechanisms that control the targeting of proteins and lipids to distinct intracellular compartments, it is becoming equally important to understand the protective mechanisms that maintain this compartmentalization in the face of challenges to membrane integrity and/or the specific biochemical identity of organelles. Here, we further explore the mechanism of ESCRT surveillance of the nuclear membranes. We interpret our data in a model where the nuclear envelope is surveilled by two principle components, the ESCRT Chm7 and the integral INM protein, Heh1. This surveillance system appears to be set up to respond directly to perturbations in the nuclear envelope barrier in a way that we suggest is agnostic as to whether the perturbation is a result of defectively formed NPCs or a mechanical (or other) disruption of the nuclear membranes.

The rationale behind this assertion is that the nuclear envelope surveillance system is itself directly established by a functioning nuclear transport system, which physically segregates Chm7 and Heh1 on either side of the nuclear envelope. For example, prior work has shown that Heh1 requires the function of the NTRs Kap-α and Kap-β1 in addition to the Ran-GTPase in order to be actively targeted to the INM through NPCs (King et al., 2006). Here, we establish that Chm7, while small enough to passively leak through the NPC diffusion barrier, is actively exported by the major export NTR, Xpo1 (Figure 1). Therefore, any perturbations that impact active nuclear transport or the diffusion barrier across the nuclear envelope would lead to increased Chm7 diffusion into the nucleus and/or a deficit in its nuclear export increasing its likelihood of meeting Heh1. While similar defects could also lead to Heh1 mistargeting and/or its diffusion into the ONM, we suspect that this would be kinetically slower than Chm7 diffusion into the nucleus as Heh1 is also bound to chromatin at the INM (Grund et al., 2008; Mekhail et al., 2008; Gonzalez et al., 2012; Yam et al., 2013; Barton et al., 2015; Schreiner et al., 2015). Indeed, it is probable that Heh1 functions in two major roles with respect to nuclear integrity: first, it provides mechanical stability to the nucleus by binding chromatin, and second, provides a binding site for Chm7 through its C-terminal WH domain. These two inter-related roles could also help to explain why both the N- and C-terminal domains of Heh1 are required to maintain viability of *apq12Δ* cells, and why *HEH1* is generally more essential in budding and fission yeasts compared to *CHM7*. This likely holds true in mammalian models as well as LEM2 is essential whereas CHMP7 is dispensable for viability (in cell culture)(Hart et al., 2015).

Once a perturbation occurs to the nuclear envelope barrier through defects in NPC assembly, loss of function of NPCs, or mechanical disruption of the nuclear membranes, Chm7 and Heh1 are able to come together. While we have so far been unsuccessful in reconstituting a direct biochemical interaction between Chm7 and Heh1 (although others have between LEM2 and CHMP7 (Gu et al., 2017)), we have nonetheless provided evidence that this interaction likely leads to Chm7 activation (Figure 4). Our data support a model in which the C-terminal WH domain of Heh1 in addition to a membrane anchor are necessary and sufficient for this activation event. As our prior data and that from Olmos *et al*. (2016), support that it is the N-terminal ESCRT-II-like domain of Chm7 (which, interestingly, is also predicted to be made up of tandem WH domains; Figure 1A) that is necessary for recruitment to the nuclear envelope, it seems likely that the binding between the Heh1-WH domain and the N-terminus of Chm7 could trigger activation by removing some form of autoinhibition, which is a common theme among ESCRT-III subunits (Zamborlini et al., 2006; Shim et al., 2007; Lata et al., 2008; Bajorek et al., 2009; Henne et al., 2012; Tang et al., 2015, 2016) that ensures polymerization in the correct compartment. Clearly the precise molecular mechanism of Chm7 activation by Heh1 will require structural insight, which is no doubt on the horizon.

Even with structural information, it will ultimately be essential to reconstitute a minimal nuclear envelope repair system that can also incorporate additional downstream components including Snf7, but also Did2, Vps2 and Vps24 (Figure 3B, C). As the latter impact the recruitment of Vps4, it is possible that the sequence of events of nuclear envelope sealing are similar to those found at ILVs during MVB formation with a fundamental difference being that a nuclear envelope hole instead of a single continuous lipid bilayer, is the likely initial “substrate” for ESCRT-III action. Indeed, in the more artificial scenarios that we present here, be it at sites of chm7_OPEN_ accumulation or Chm7 in *vps4Δ* cells (Figures 6 and 7), there are obvious parallels between the morphologies observed at the INM and those at the plasma membrane (von Schwedler et al., 2003; Hanson et al., 2008; Morita et al., 2011; Cashikar et al., 2014; Jackson et al., 2017), and at endosomes (Adell et al., 2014; Wenzel et al., 2018) in the context of mutants that stall or inhibit membrane scission. They also resemble more physiological circumstances like in the ILVs of *A. thaliana* (Buono et al., 2017) and *C. elegans* (Frankel et al., 2017), which resemble beads on a string. The network of INM evaginations also resemble the fenestrated membrane cisternae formed in the ER-derived unconventional secretion pathway, CUPS (Curwin et al., 2016).

Additionally, the size of the necks at the base of the INM evaginations and nuclear envelope herniations is suggestive of the presence of a spiraling polymer analogous to those observed in vitro and in vivo (McCullough et al., 2018), supporting that these necks are likely stabilized by ESCRTs. Thus, while we acknowledge that such INM evaginations might not be a physiological event in the nuclear envelope sealing process, it remains tempting to speculate that they might be in the context of proposed mechanisms of nuclear egress be it Mega-RNPs (Speese et al., 2012; Jokhi et al., 2013), viruses (Lee et al., 2012, 2016; Arii et al., 2018), or in nucleophagy mechanisms (Roberts et al., 2003; Dou et al., 2015; Mochida et al., 2015; Mostofa et al., 2018) that so-far remain obscure but would require a membrane scission step. Interestingly, recent work suggests that herpes virus nuclear egress requires ESCRTs (Arii et al., 2018), including a role in controlling interesting INM extensions into the nucleus; such intranuclear membrane might also be relevant in cell types that have so-called “nucleoplasmic reticulum” (Malhas et al., 2011). The biogenesis and function of nucleoplasmic reticulum remain enigmatic but our observations of intranuclear fenestrated membrane emanating from the INM might suggest a yet-to-be discovered role for Chm7 and Heh1 in forming such structures as well.

That NPC biogenesis during interphase also proceeds through an INM evagination (Otsuka et al., 2016) step raises the possibility that ESCRTs might also directly contribute to this process. However, while this remains a compelling hypothesis, our data continue to argue against this possibility. For example, the herniations observed in *apq12Δ* and *nup116Δ* cells are ostensibly products of either defective INM-ONM fusion during NPC biogenesis (which is downstream of INM evagination), and/or, a triggering of a NPC assembly surveillance mechanism (Thaller and Lusk, 2018). In the former case, an expectation would be that deletion of *CHM7* would prevent the formation of nuclear envelope herniations, which clearly remain present in *apq12Δchm7Δ* strains (Figure 8C, D). In the latter case, the expectation would be that the nuclear envelope herniations would be unsealed without *CHM7*. It is highly unlikely, however, that we would be able to capture a herniation that is “open” as these membranes would be inherently unstable. Indeed, that we observe often-dramatic openings in the nuclear envelope in *apq12Δchm7Δ* cells (Figure 8D and Figure 8 - figure supplement 2), suggest that the herniations themselves might be prone to rupture. It follows then that like in mammalian cells where much larger nuclear envelope herniations are precursors to nuclear rupture (de Vos et al., 2011; Vargas et al., 2012; Hatch et al., 2013; Denais et al., 2016; Hatch and Hetzer, 2016; Raab et al., 2016), these smaller NPC-assembly associated herniations might also impact nuclear envelope integrity through mechanisms that remain to be fully defined. In either case, it reinforces the concept that the assembly of NPCs can be perilous, and it will be important to consider this possibility when interpreting the underlying pathology of human diseases that are associated with defects in NPC function or assembly, for example, DYT1 early-onset dystonia (Laudermilch et al., 2016; Pappas et al., 2018) or Steroid Resistant Nephrotic Syndrome (Miyake et al., 2015; Braun et al., 2016, 2018).

Interpreting the pathology of genetic diseases associated with nuclear envelope malfunction will also benefit from a more mechanistic understanding of how the activation of this surveillance pathway leads to nuclear membrane sealing. Two of the more interesting observations of the EM tomographic analyses are the generation of intranuclear membrane invaginations and the proximity of ER sheets and vesicles at the ONM sites of nuclear envelope herniations (Figure 6, Figure 6 - figure supplement 1, Figure 7, figure 7 - figure supplement 1). Both of these observations suggest that nuclear envelope sealing might be coupled to the delivery of new membrane at sites of rupture. Such a mechanism might be analogous to the reformation of the nuclear envelope at the end of mitosis in mammalian cells, which requires ESCRTs and the recruitment of ER membranes to the chromatin surface (Olmos et al., 2015, 2016; Vietri et al., 2015). It seems plausible that sealing holes in the nuclear envelope (particularly those larger than a nuclear pore) would require the local delivery of membranes. Similarly, NPC quality control mechanisms that suggest the sealing of defective NPCs would likely require some form of expansion of the pore membrane, as has been proposed (Wente and Blobel, 1993). Whether such membrane is derived from new synthesis or from the mobilization of existing stores remains to be explored. Of note, recent work supports that even the INM may be metabolically active and have the capacity to generate new lipid locally (Romanauska and Köhler, 2018). Further, close examination of samples reveals that the morphology of the INM extensions are both sheet-like and tubular in nature - it is not immediately obvious how such membrane connections could be established but they closely resemble the ‘normal’ continuities between the ONM and the broader ER network (West et al., 2011); together these data suggest that ER-shaping proteins and lipid synthesis pathways might play an important role in contributing to nuclear envelope sealing - topics for the future.

Lastly, any ESCRT-mediated nuclear envelope sealing mechanism will need to incorporate a potential role for NTRs. The sequence overlap of the potential MIM domain and one of the Chm7 NESs (Figure 1A, Figure 1 – figure supplement 1A) raises the possibility that Xpo1 might directly impact Chm7 function at the nuclear envelope, perhaps even playing a direct role in polymer formation. Likewise, Heh1 is synthesized and inserted into ER membranes before it is targeted to the INM by NTRs. As exposure of Heh1 to the cytosol is sufficient to induce Chm7 recruitment to ER membranes, do NTRs inhibit Chm7 recruitment in this compartment? The latter concept points to a similar level of regulation that must be imposed in mammalian cells during mitotic nuclear envelope breakdown in which LEM2 would be exposed to CHMP7. Whether NTRs also bind to LEM2 in mammalian cells remains an outstanding question but one that is supported by the presence of NLS-like sequences in its N-terminus (Kralt et al., 2015). Regardless, recent work is already suggesting additional regulatory proteins like Lgd/CC2D1B that control the spatiotemporal timing of CHMP4B and CHMP7 activity in mammalian cells (Ventimiglia et al., 2018). These, and likely other yet-to-be defined factors promise new insights into this emerging mechanism of organelle surveillance.

## Acknowledgements

We thank members of the Lusk, Beck and King laboratories for critical input in addition to M. King for comments on the manuscript. We also acknowledge invaluable assistance from Z. Hakhverdyan and M. Rout for affinity purifications and the Yale Keck Proteomics facility, M. Graham and X. Liu for help with EM and generous support by EMBL’s electron microscopy core facility and Y. Schwab. This work was supported by the NIH, GM105672 to C.P.L. and D.J.T. was also funded by 5T32GM007223 and a short term EMBO fellowship 6885. M.A was funded by an EMBO a long term fellowship (ALTF-1389-2016).

---

**Figure 1 – figure supplement 1:**
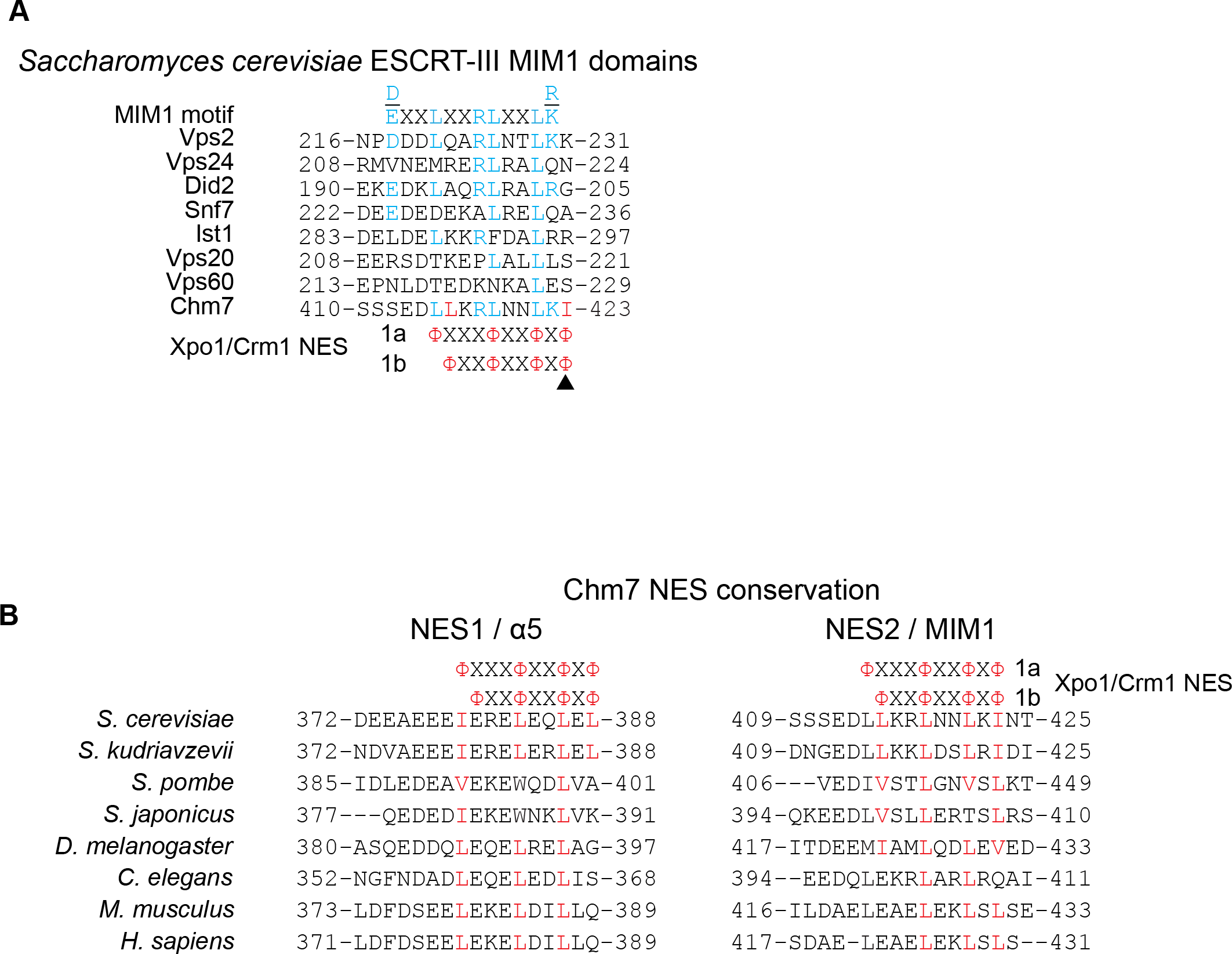
Chm7 can diffuse across the NPC but is actively exported by Xpo1. **A**) Amino acid sequences of *S. cerevisiae* ESCRT-III MIM1 domains aligned with class 1a and 1b Xpo1/Crm1 NESs. Blue coloring reflects alignment with the MIM1 consensus (at top) and red coloring indicates additional hydrophobic amino acid residues in the putative Chm7 MIM1 domain that align with both the class 1a and 1b NESs. **B**) Alignment of predicted NES1 and NES2 from Chm7 with the analogous sequence of the indicated species. Red coloring are hydrophobic residues that align with type 1a and 1b NES consensus sequences.

**Figure 3 – figure supplement 1:**
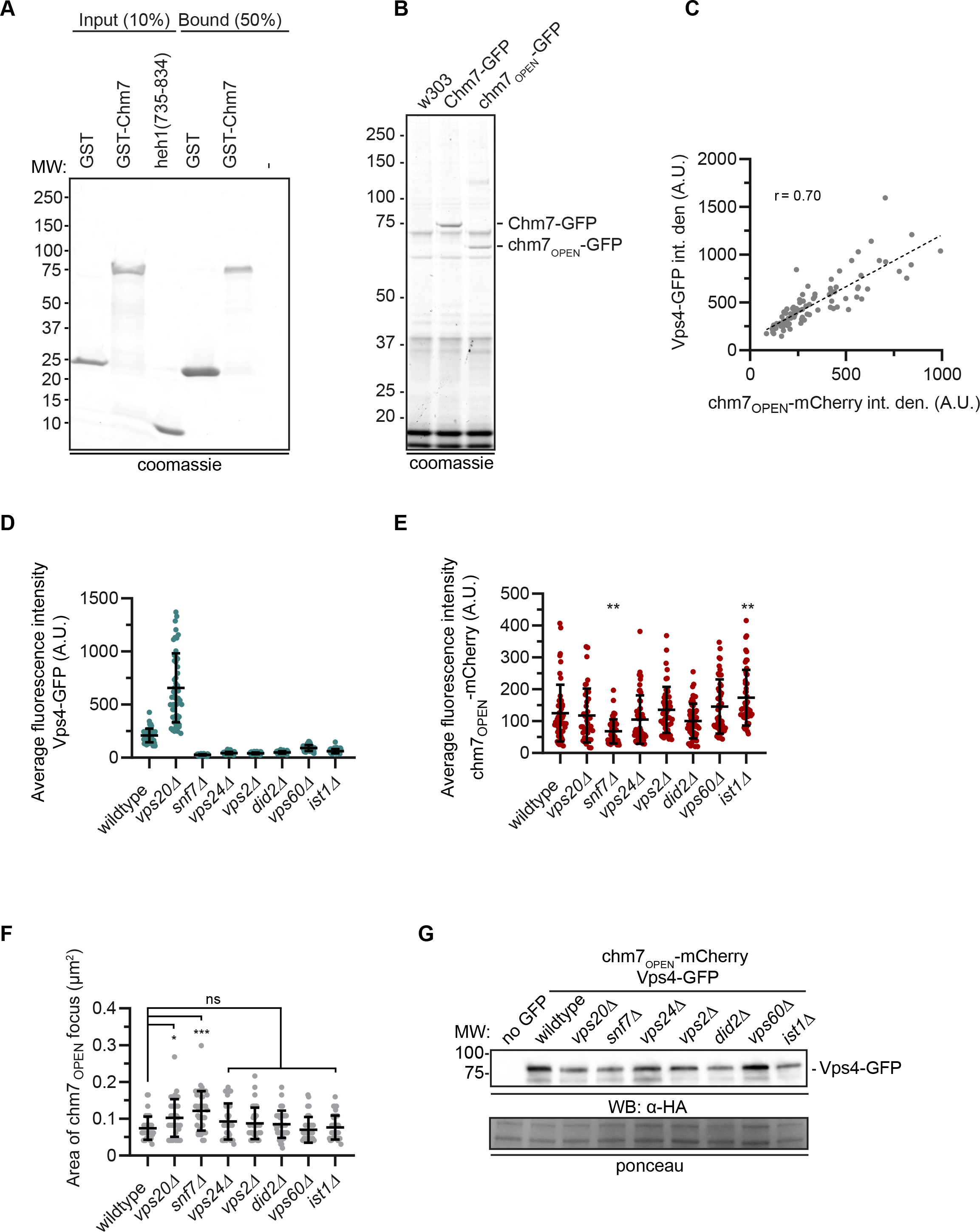
Chm7 binds to Heh1, Nur1 and downstream ESCRTs required for Vps4 recruitment. **A**) Recombinant purified Heh1 WH domain and GST-Chm7 fail to interact within an in vitro solution binding assay. A coomassie-stained SDS-PAGE gel showing input and bound fractions of a solution binding experiment where GST or GST-Chm7 were incubated with the Heh1 WH domain (heh1(735-834)) and immobilized on GT-beads. The position of MW markers are indicated at left. **B**) Anti-GFP magnetic beads were incubated with cell extracts derived from wildtype (W303) or cells expressing endogenous levels of Chm7-GFP or chm7_OPEN_-GFP. Proteins bound to beads were eluted, separated by SDS-PAGE and either sent for MS/MS peptide identification or stained with Coomassie as shown. Position of molecular weight (MW) markers are shown at left. **C**) Correlation of the total fluorescence (integrated density) in arbitrary units (A.U.) of chm7_OPEN_-mCherry and Vps4-GFP co-localized at the nuclear envelope. Linear regression calculated from 100 chm7_OPEN_ foci pooled from three independent replicates; *r* is the linear correlation (Pearson’s) coefficient. **D**) Plot of average fluorescence intensity (in arbitrary units; A.U.) of Vps4-GFP co-localized with chm7_OPEN_-mCherry foci in the indicated genetic backgrounds. Error bars represent SD of the mean from > 50 foci pooled from three independent replicates. **E**) Plot of average fluorescence intensity (in arbitrary units) of chm7_OPEN_-mCherry foci in the indicated genetic backgrounds. Error bars represent SD of the mean from >50 foci pooled from three independent replicates. *P*-values from one-way ANOVA with Dunnett’s correction where ** *P* ≤ 0.01. **F**) Plot of the area encompassed by the chm7_OPEN_-GFP focus in the indicated genetic backgrounds. Error bars represent SD of the mean from > 50 foci pooled from three independent replicates. *P*-values from one-way ANOVA with Dunnett’s correction where ns is *P* ≥ 0.05, * *P* ≤ 0.05, and *** *P* ≤ 0.001. **G**) Western blot to assess the total levels of Vps4-GFP in the indicated genetic backgrounds. Anti-HA-labeling detected by HRP-conjugated secondary antibodies and ECL. Ponceau-stain used to assess total protein loads. Position of MW markers at left.

**Figure 4 – figure supplement 1:**
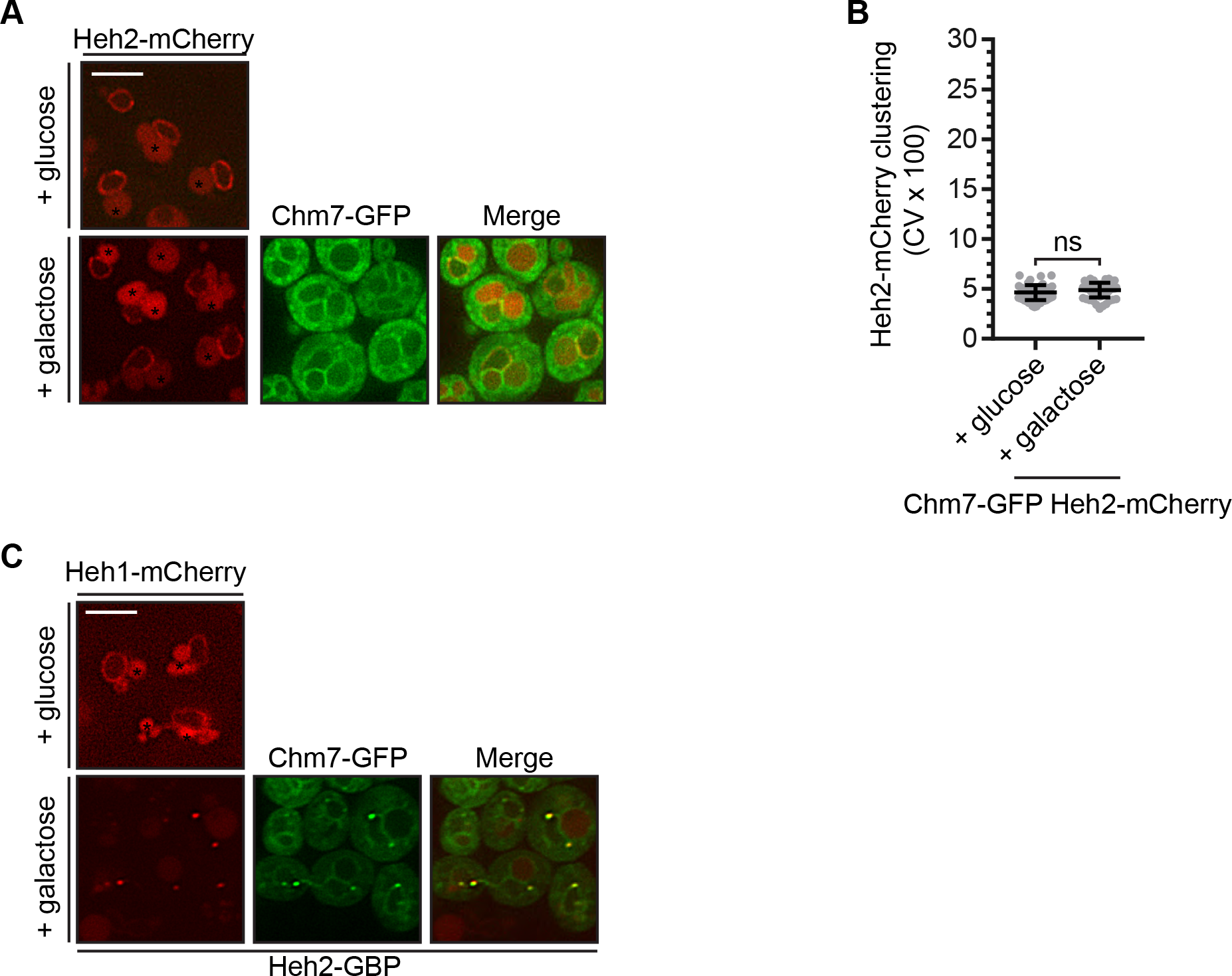
Heh1 is required to induce the focal accumulation of Chm7. **A**) Overexpression of Chm7-GFP does not impact the distribution of Heh2-mCherry. Deconvolved fluorescence micrographs of Heh2-mCherry and Chm7-GFP under conditions where Chm7-GFP produced (galactose) or repressed (glucose). Merge of green and red fluorescence also shown. Asterisks denote vacuolar autofluorescence. Scale bar 5 μm. **B**) Plot of the CV of Heh2-mCherry fluorescence along the nuclear envelope from images and conditions in (A). Error bars represent SD of the mean from three independent replicates of > 50 cells. *P*-values from Student’s t-test where ns is *P* ≥ 0.05. **C**) Overexpression of Chm7-GFP in the context of Heh2-GBP leads to the accumulation of Heh1-mCherry in Chm7-GFP foci at the nuclear envelope. Deconvolved fluorescence micrographs of Heh1-mCherry and Chm7-GFP (in a strain also expressing Heh2-GBP) under conditions where Chm7-GFP is produced (galactose) or repressed (glucose). Merge of green and red fluorescence also shown. Asterisks denote vacuolar autofluorescence. Scale bar 5 μm.

**Figure 5 – figure supplement 1:**
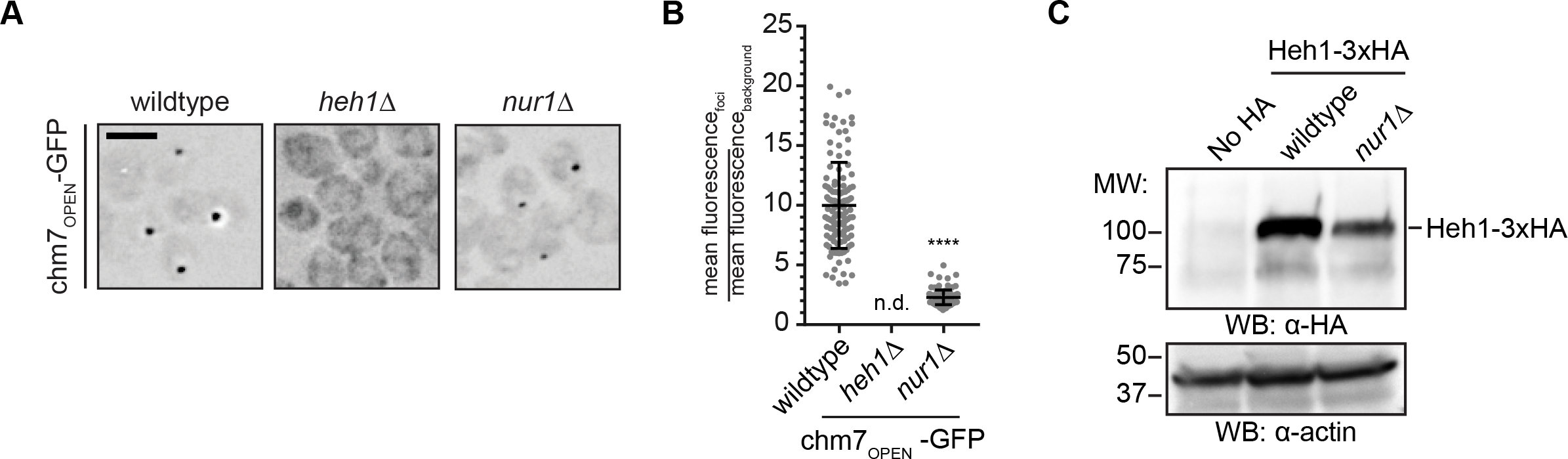
The Heh1 WH domain and a transmembrane anchor are necessary and sufficient for Chm7 activation. **A**) Deletion of *NUR1* impacts chm7_OPEN_-GFP accumulation at the nuclear envelope. Deconvolved fluorescence micrographs (inverted) of chm7_OPEN_-GFP in the indicated strains. **B**) Plot of the level of enrichment of chm7_OPEN_-GFP at nuclear envelope foci in the indicated strains reflected by the ratio of its mean fluorescence at the nuclear envelope focus over background fluorescence. Error bars represent SD of the mean of >100 foci pooled from three independent replicates. **C**) Heh1 levels are diminished in *nur1Δ* cells. Western blot of the total levels of a Heh1-3xHA fusion (produced from endogenous promoter) in the indicated strains. Heh1-3xHA and actin, which is used as a total protein load reference, are detected using either anti-HA antibodies or anti-actin antibodies followed by HRP-conjugated secondary antibodies and ECL. Position of molecular weight (MW) standards at left.

**Figure 6 – figure supplement 1:**
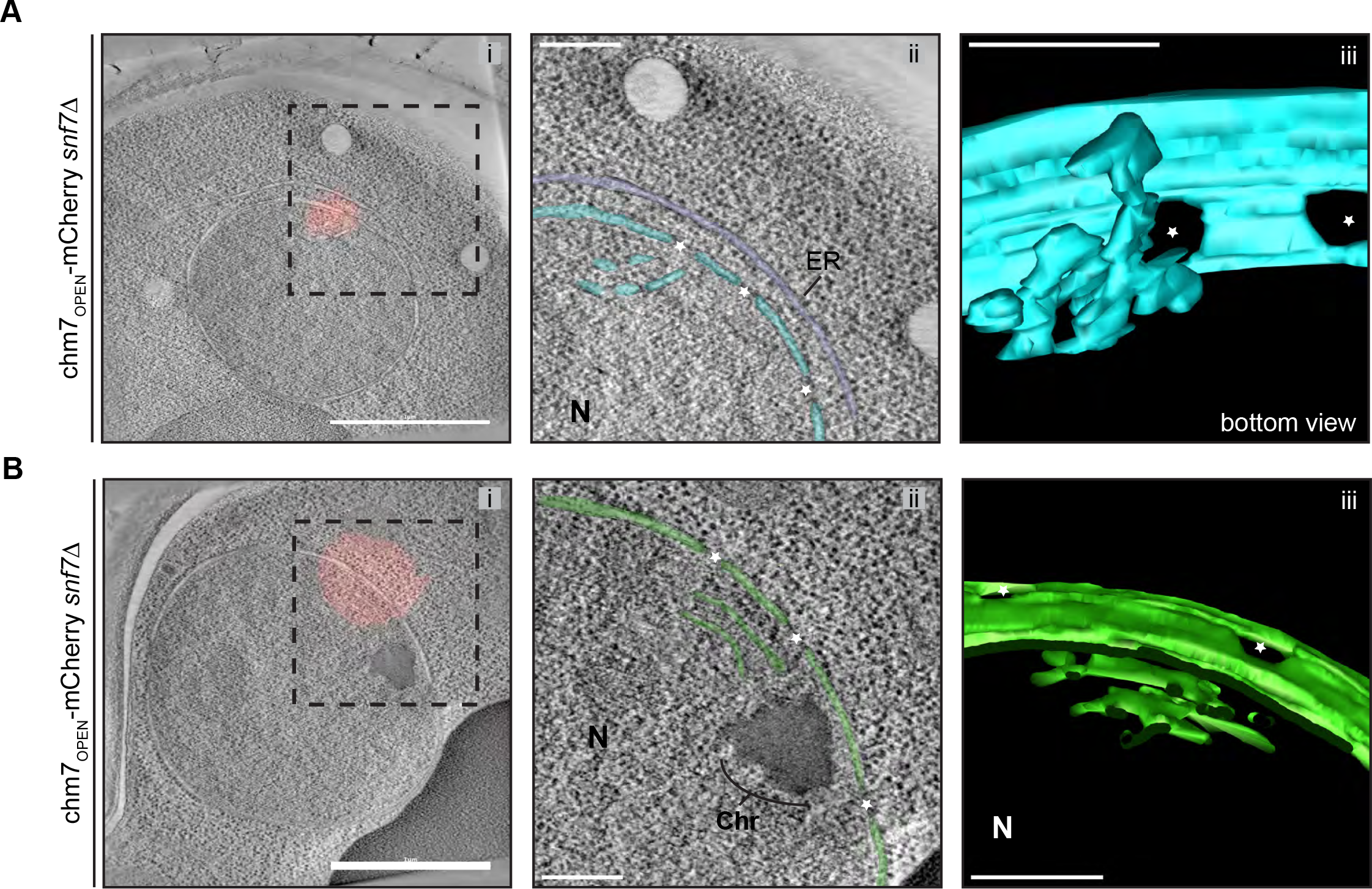
chm7_OPEN_ associates with a network of intranuclear fenestrated cisterna often below nuclear envelope herniations. **A and B**) *SNF7* is dispensable for the formation of an intranuclear fenestrated membrane network in the context of chm7_OPEN_. *i*) Correlative light and electron microscopy of ~220 nm thick sections was used to examine the morphology of the nuclear envelope at sites of chm7_OPEN_-mCherry accumulation in *snf7Δ* cells. *i.* Overlay of fluorescent (red) and electron micrographs; boxed region is magnified in *ii. ii.* Tomographic slices where nuclear envelope membranes are colored blue or green with ER in purple. *iii*. Views of 3D models. Stars are nuclear pores, Chr is chromatin and N is nucleus. Scale bars in *i* are 1 μm and scale bars in *ii*, *iii* are 250 nm.

**Figure 8 – figure supplement 1:**
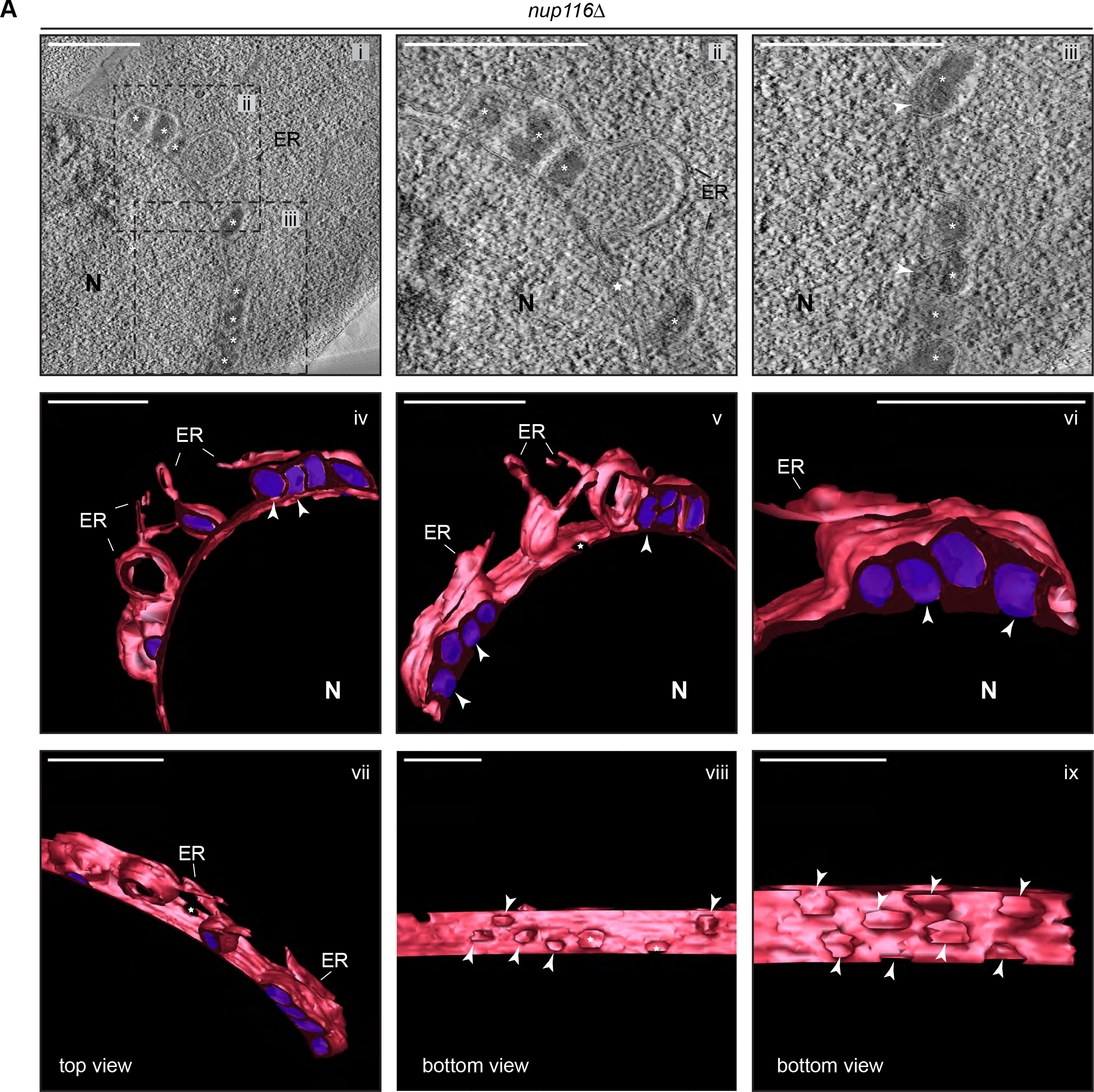
*CHM7* is required to maintain the integrity of the nuclear membranes in the context of nucleoporin-associated herniations. **A)** Tomographic slices from 300 nm sections of *nup116Δ* cell nuclear envelope herniations. *i-iii*, Tomographic slice with boxed regions magnified in *ii* and *iii*. *iv-ix*. Different persepectives of 3D model with nuclear envelope and ER membranes in red and herniation densities (marked by asterisks in i-iii) are colored purple. White arrowheads point out the bases of the herniations and stars are nuclear pores. Scale bars are 250 nm.

**Figure 8 – figure supplement 2:**
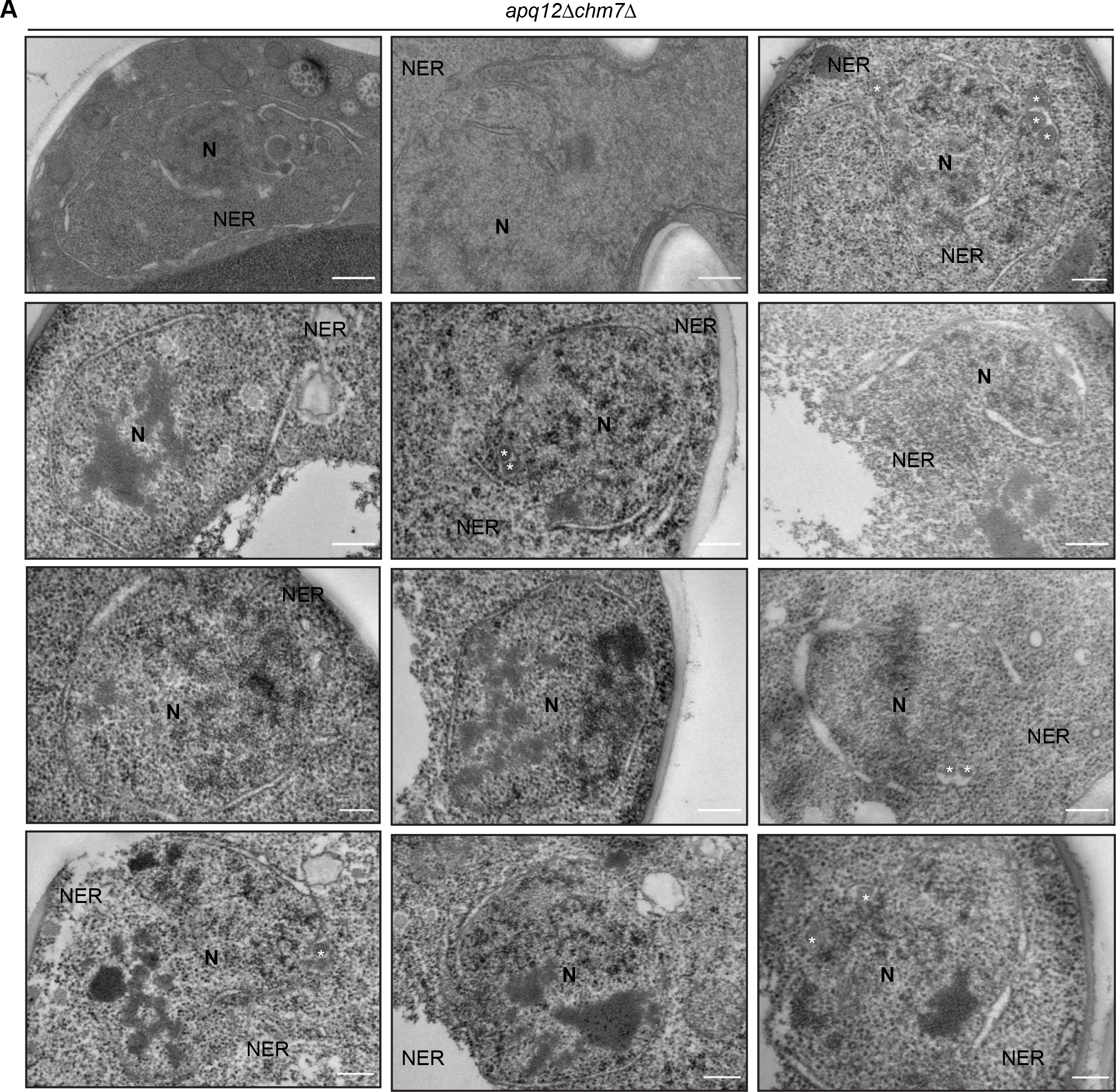
*CHM7* is required to maintain the integrity of the nuclear membranes in the context of nucleoporin-associated herniations. **A)** Electron micrographs (from thin sections) showing nuclear envelope ruptures (NER) in *apq12Δchm7Δ* cells after 2 hours at 37oC. Asterisks indicate herniations. Scale bars are 250 nm.

**Figure 8 – figure supplement 3:**
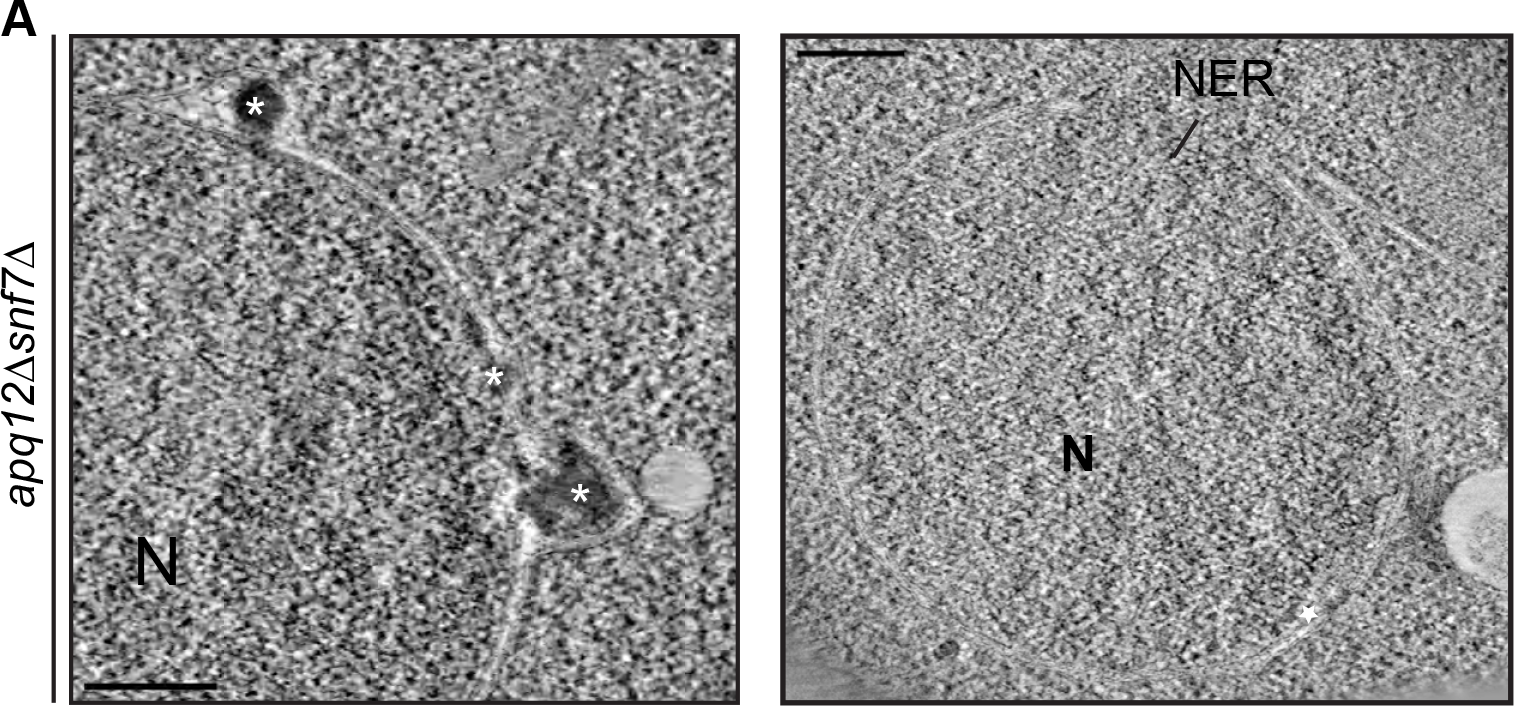
*CHM7* is required to maintain the integrity of the nuclear membranes in the context of nucleoporin-associated herniations. **A)** Electron micrographs (from thin sections) showing nuclear envelope ruptures (NER) and herniations (asterisks) in *apq12Δsnf7Δ* cells after 2 hours at 37°C. Scale bars are 250 nm

## Video Legends

**Video 1: Clustering of Heh2-GBP-mCherry in FETA assay requires Heh1.** Related to Figure 4. A timelapse series of fluorescence images acquired at 10 minute intervals of overexpressed Chm7-GFP and Heh2-GBP-mCherry in wildtype and *heh1Δ* cells. Green, red and merge shown. Timestamp shows elapsed time after galactose induction of Chm7-GFP expression. Scale bar is 2 μm.

**Video 2: Nuclear envelope morphology at sites of chm7_OPEN_-GFP accumulation.** Related to Figure 6A. Video showing full tomogram and 3D model from a nuclear envelope region of chm7_OPEN_-GFP accumulation. Scale bar is 250 nm.

**Video 3: Nuclear envelope morphology at sites of chm7_OPEN_-GFP accumulation.** Related to Figure 6B. Video showing full tomogram and 3D model from a nuclear envelope region of chm7_OPEN_-GFP accumulation. Scale bar is 250 nm.

**Video 4: Nuclear envelope morphology at sites of chm7_OPEN_-GFP accumulation in the absence of *SNF7*.** Related to Figure 7A. Video showing full tomogram and 3D model from a nuclear envelope region of chm7_OPEN_-mCherry accumulation. Scale bar is 250 nm.

**Video 5: Morphology of nuclear envelope herniations in *apq12Δ* cells.** Related to Figure 8C. Video showing a tomogram and 3D model of the nuclear envelope in *apq12Δ* cells. Scale bar is 250 nm.

**Video 6: Morphology of nuclear envelope herniations in *nup116Δ* cells.** Related to Figure 8 – figure supplement 1A. Video showing a tomogram and 3D model of the nuclear envelope in *nup116Δ* cells. Scale bar is 250 nm

## MATERIALS AND METHODS

### Yeast strains and growth conditions

All strains used in this study are from a W303 parent; their derivation and genotypes are listed in **Table 1**. Fluorescent protein tagging and gene deletions were generated using a PCR-based integration approach using the pFA6a plasmid series (**Table 2**) as templates (Longtine et al., 1998; Van Driessche et al., 2005). Standard yeast protocols for transformation, mating, sporulation, chromosomal DNA isolation and tetrad dissection were followed (Amberg et al., 2005).

Cells were grown to mid-log phase in YPA (1% Bacto yeast extract (BD), 2% Bacto peptone (BD), 0.025% adenine hemi-sulfate (Sigma)) or complete synthetic medium (CSM) supplemented with 2% raffinose (R; BD), 2% D-galactose (G; Alfa Aesar) or 2% D-glucose (D; Sigma) as indicated.

To compare relative growth rates of *heh1Δapq12Δ* strains expressing *HEH1* alleles (DTCPL1498, DTCPL1517, DTCPL1581, DTCPL1519, DTCPL1520) roughly equivalent cell numbers from overnight cultures grown in in YPAR were spotted in 10-fold serial dilutions onto YPG to induce expression of Heh1 or indicated truncations and imaged after 36 hours at RT.

### Leptomycin B and BAPTA-AM treatments

To test the impact of inhibiting nuclear export on the steady state distribution Chm7-GFP, Chm7-MGM4, chm7_OPEN_-GFP, GFP, NES2_CHM7_-GFP, or NES1-NES2_CHM7_-GFP, we used KWY175 (a gift from B. Montpetit and Karsten Weis) in which the genomic deletion of *XPO1* is covered with pRS413 expressing the *xpo1-T539C* allele that confers sensitivity to Leptomycin B (LMB). These strains were grown in YPAR and galactose (final concentration of 1%) was added to the growth medium to induce the expression of the GFP fusion proteins for 2 hours before the addition of 2% D-glucose to repress protein production. Cultures were then treated with 50 ng/mL LMB dissolved in 7:3 MeOH:H_2_O solution (Sigma) for 45 minutes alongside a control of the equivalent volume of MeOH before imaging.

To test if Ca^2+^ plays a role in the physiological recruitment of Chm7 to the nuclear envelope, *apq12Δ* cells expressing Chm7-GFP (DTCPL567) were cultured overnight at 30°C, diluted to an OD_600_ of 0.2 and grown for an additional 2 hours at RT. Cells were treated with either 25 μM (Li et al., 2011) of the cell permeable calcium chelater BAPTA-AM (Tocris Bioscience) dissolved in DMSO or DMSO alone for 30 minutes followed by a 45 minute incubation at either RT or 37°C before imaging.

### Plasmids

All plasmids are listed in **Table 2.**

To generate pRS406-ADH1-GFP, the GFP coding sequence was amplified by PCR and inserted into pRS406-ADH1 (p406ADH1 was a gift from Nicolas Buchler & Fred Cross-Addgene plasmid # 15974; http://n2t.net/addgene:15974; RRID:Addgene_15974)) using *Eco*RI and *Hind*III restriction sites.

To generate pCHM7-MGM4, the *CHM7* ORF was amplified by PCR with *Cla*I restriction sites and subcloned in pPP004 linearized with *Cla*I (gift of L. Veenhoff; (Popken et al., 2015)), placing *CHM7* in between the first MBP gene and the GFP. To generate pRS406-ADH1-NES2-GFP, complimentary 4 nmol Ultramers (IDT) were designed to code for amino acids 409-424 of *CHM7* with overhangs that would be generated with *Xba*I or *Bam*HI. Ultramers were annealed by heating to 95°C for 5 minutes in Taq polymerase PCR buffer (Invitrogen) and allowing them to slowly cool to RT. Annealed primers were then ligated using T4 ligase (Invitrogen) into pRS406-ADH1-GFP linearized with *Xba*I*/Bam*HI (New England Biolabs).

To generate pRS406-ADH1-NES1-NES2_CHM7_-GFP, the sequence encoding amino acids 370-429 of *CHM7* was amplified with oligonucleotide primers containing the *Xba*I or *Bam*HI restriction sites. The PCR product was digested with *Xba*I*/Bam*HI and gel purified (Qiagen) before ligation with T4 ligase (Invitrogen) into pRS406-ADH1-GFP linearized with *Xba*I*/Bam*HI.

Gibson Assembly (New England BioLabs) was used to generate pFA6a-3xHA-GFP-his3MX6 for functional tagging of Vps4(Adell et al., 2017). The 3xHA epitope was PCR-amplified from a pFA6a-3xHA-his3MX6 (Longtine et al. 1998) plasmid using Q5 DNA polymerase (New England BioLabs) and assembled into pFA6a-GFP-hisMX6 (Longtine et al. 1998) digested with *Sal*I and *Pac*I (New England BioLabs).

### Western blotting

For whole cell protein extracts, approximately 2 OD_600_ of cells in mid log phase were collected by centrifugation, washed in 1 mM EDTA, pelleted again and resuspended/lysed in 2 M NaOH for 10 minutes on ice. Proteins were precipitated by the addition of 50% TCA (Trichloroacetic acid) for 20 minutes on ice and then collected by centrifugation. The precipitated proteins were washed in ice-cold acetone, air dried and then resuspended in SDS-PAGE sample buffer. Samples were then denatured by at 95°C for 5 minutes. Denatured proteins were separated on precast SDS-PAGE, 4-20% gradient gels (BioRad) and transferred to 0.2 μm nitrocellulose membranes (BioRad). Relative protein loading was visualized using Ponceau S Solution (Sigma). Membranes were subsequently washed in TBST and blocked for 1 hour in 5% skim milk in TBST at RT. Membranes were then incubated with HRP-conjugated anti-HA (Roche 3F10), or anti-actin (mAbcam 8224) diluted in TBST. Primary antibodies were detected directly with ECL (ThermoFisher) or with anti-rabbit HRP-conjugated secondary antibodies, followed by ECL and visualized using a VersaDoc Imaging System (Bio-Rad).

### Fluorescence microscopy

With the exception of the correlative light electron microscopy experiments described below, all fluorescence micrographs were acquired using a DeltaVision microsope (Applied Precision/GE Healthcare) fitted with a 100×, 1.4 NA objective (Olympus). Images were taken using a CoolSnapHQ^2^ CCD camera (Photometrics), with the exception of those in Figure 3 and Figure 4B which were acquired using a Evolve EMCCD camera (Photometrics).

For timecourse assessment of Heh2-GBP-mCherry clustering in FETA assay in Figure 4B, cells were imaged in microfluidic plates (Y04C/CellASIC) in the ONIX microfluidic platform (CellASIC). Cell were loaded into the microfluidic chamber in CSM with 2% raffinose. CSM with 2% galactose was perfused into the microfluidic chamber at 0.25 psi for the course of the experiment. Z-stacks (0.4 μm sections) were acquired for 90 minutes at 10 minute intervals.

### Image processing and analysis

All presented fluorescent micrographs were deconvolved using an iterative algorithm in softWoRx (6.5.1; Applied Precision GE Healthcare). Unprocessed images after background subtraction were used for quantification of fluorescence intensities. Assessment of Heh2-GBP-mCherry or Heh2-mCherry clustering was quantified by calculating the coefficient of variation (SD/mean × 100) of fluorescence of individual nuclear envelopes(Fernandez-Martinez et al., 2012). A 4 pixel wide, freehand line was traced over the entire nuclear envelope in a mid-plane section using FIJI/ImageJ (Schindelin et al., 2012) and the mean fluorescence contained in the traced area was measured.

To correlate the fluorescence intensity of co-localized Vps4-GFP and chm7_OPEN_-mCherry, the integrated density of Vps4-GFP and Chm7_OPEN_-mCherry was measured and plotted on a correlation curve. The linear correlation coefficient (Pearson coefficient, r) was calculated in Prism (GraphPad 8.0). Similarly, quantification of the integrated density and average fluorescence intensity of Vps4-GFP and chm7_OPEN_-mCherry were measured by selecting a region of interest (ROI) around the chm7_OPEN_-mCherry signal and measuring average fluorescence intensity in both mCherry and GFP channels.

To measure relative nuclear exclusion of GFP, NES2_CHM7_-GFP, and NES1-NES2_CHM7_-GFP constructs at steady state, a 3.75 μm line was traced across each nucleus (encompassing cytoplasm both times the line crosses the nuclear envelope border) as determined from the dsRed-HDEL localization. GFP fluorescence was measured using the Plot Profile function in FIJI/ImageJ (Schindelin et al., 2012). Traces were normalized to the maximum value measured within each trace before averaging.

### Statistical Analyses

Graphs and statistical analyses were generated using Prism (GraphPad 8.0). *P*-values in all graphs were generated with tests as indicated in figure legends and are represented as follows: ns, *P* > 0.05; * *P* ≤ 0.05; ** *P* ≤ 0.01 *** *P* ≤ 0.001, **** *P* ≤ 0.0001. All error bars represent the standard deviation from the mean. Scatter plots of spectral counts from MS/MS analysis for Figure 3A, B, and C were generated using Excel (Microsoft).

### Nuclear export signal prediction

Xpo1/Crm1 NES sequences were predicted using LocNES (Xu et al., 2015) with default threshold settings.

### Recombinant protein binding experiments

GST, GST-Chm7 and GST-heh1(735-834) proteins were recombinantly produced and purified as previously described (Webster et al., 2016) in lysis buffer (50 mM Tris pH 7.4, 500 mM NaCl, 2 mM MgCl_2_, 2 mM CaCl_2_,10% glycerol, 0.5% NP-40, 1 mM DTT, complete protease inhibitors (Roche)). The soluble fraction was incubated with glutathione sepharose (GT) beads for 1h at 4°C for binding. The GT beads were collected by centrifugation and washed thrice with lysis buffer. GST and GST-Chm7 proteins were eluted from GT beads by 10 mM reduced glutathione and dialyzed in lysis buffer. heh1(735-834) was cleaved off from GT beads by incubating with HRV3C protease (Thermo Scientific) at 4°C overnight. For binding experiment, heh1(735-834) was incubated with GST and GST-Chm7 in a dialysis cassette (3.5K Slide-A-Lyzer, Thermo Scientific) and the binding reaction was dialyzed overnight in a binding buffer (50 mM Tris pH 7.4, 150 mM NaCl, 2 mM MgCl_2_, 2 mM CaCl_2_,10% glycerol, 0.5% NP-40, 1 mM DTT). The reaction was collected, incubated with GT beads for 1 h at 4°C, washed thrice with binding buffer and eluted in 2X Laemmli sample-buffer. Proteins were resolved on a SDS-PAGE gel and visualized by SimplyBlue Safe-Stain (Invitrogen).

### Immunoaffinity purification

*S. cerevisiae* cells expressing Chm7-GFP (DTCPL81), Chm7-GFP *vps4Δpom152Δ* (DTCPL133), chm7_OPEN_-GFP (DTCPL413) were grown to log phase in YPD at 30°C and collected by centrifugation. Cells were washed once with ice-cold water, collected by centrifugation and resuspended in a small volume of freezing buffer (20 mM HEPES, pH 7.4, 1.2% polyvinylpyrrolidone and protease inhibitor (Sigma)(Oeffinger et al., 2007) and flash frozen in liquid nitrogen. The frozen yeast pellets were pulverized in a Retsch MM400 mixer mill for 6 times at 30 Hz for 3 minutes. For immunoaffinity purification, 200 mg of frozen, ground yeast powder was solubilized in 4 volumes of homogenization buffer (400 mM trisodium citrate, pH 8, 0.5% n-Dodecyl β-D-maltoside) and protease inhibitor cocktail (Roche). The soluble fraction was incubated with 10 μl magnetic beads (Dynabeads, M-270 Epoxy, Invitrogen) slurry coated with GFP-nanobody for 1 hour at 4°C (Cristea et al., 2005; LaCava et al., 2016). The beads were collected on a magnetic rack and washed three times with 500 μl homogenization buffer. Bound proteins were eluted by incubating the beads in 20 μl 1X NuPAGE LDS (lithium dodecyl sulfate) sample buffer (Invitrogen) at 70°C for 10 minutes. Eluates were separated on a magnetic rack and further incubated with 50 mM DTT at 70°C for 10 minutes. The eluates were run on a 4-12% NuPAGE gel (Novex) until the dye front just entered the gel. The gels were stained with Imperial protein stain (Thermo Scientific) and protein bands were excised for MS analysis.

### Mass Spectrometry and analysis

MS/MS was performed at the Yale Keck Proteomics facility. Excised bands described above were transferred to clean 1.5 mL Eppendorf tubes and digested with trypsin. Subsequently, chromatographic separation of peptides was done using a Waters nanoACQUITY ultra high pressure liquid chromatograph (UPLC), and peptides were detected on a Waters/Micromass AB QSTAR Elite. Analysis of MS/MS peptide results was completed using Scaffold 4.8.7 (Proteome Software Inc.). Peptides were identified by SEQUEST and Mascot using X!Tandem (Craig and Beavis, 2003; Searle et al., 2008) and validated using PeptideProphet (Keller et al., 2002; Nesvizhskii et al., 2003) within Scaffold software (Proteome Software Inc.). Proteins were identified by comparison with SwissProt database where peptide identifications required ≥ 2 peptides from each replicate and ≥ 95.0% probability of correct identification to be included in analysis. Quantitative analysis to determine significance of enrichment between samples was done with total spectral counts from two replicates using Fischer’s exact test with a significance threshold *P* < 0.05 (Figure 3A, C), or on presence/absence from 1 replicate (Figure 3B).

### Correlative fluorescence and electron tomography

Correlated fluorescence and electron microscopy were conducted as previously described (Kukulski et al., 2012; Curwin et al., 2016). In brief, yeast cells were high pressure frozen (HPM010, AbraFluid), freeze substituted (EM-AFS2, Leica) with 0.1% uranyl acetate in acetone and infiltrated with Lowicryl. 300 nm sections were cut with a microtome (EM UC7, Leica) and picked up on carbon coated 200 mesh copper grids. 50 nm TretraSpeck fluorescent microspheres (fluorescence and electron dense fiducials, Life technologies, Carlsbad, CA) were added to the grid for correlation. Grids used for Figure 6A,B, Figure 6 - figure supplement 1A,B, Figure 8C, Figure 8 - figure supplement 1A, and Figure 8 - figure supplement 3A, were poststained with lead citrate to increase contrast. In all cases, 15 nm protein A-coupled gold beads were adsorbed on both sides of each grid and used as fiducial markers for overlaying high and low magnification tomograms. 60° to −60° tilt series were acquired on a Technai F30 (Thermofisher, FEI) at 300 kV with Serial-EM (Mastronarde, 2005) at 20000× and either 3900× or 4700× to facilitate ease of correlation with TetraSpeck fiducials.

To perform CLEM, fluorescence images were acquired of the EM grids on images a Nikon TI-E (Figure 6A, B) with sCMOS PCO edge 4.2 CL camera and solid state illumination, or an Olympus IX81 with MT20 (Olympus) lamp and CCD (Orca-ER; Hamamatsu Photonics) (Figure 7A, B, Figure 6 - figure supplement 1A, B). To distinguish protein fluorescence signal from fluorescent fiducials, for each field of view/grid four channels were acquired (GFP, mCherry/RFP, Cy5, and brightfield).

Acquired images were further processed in FIJI using the Extended Depth of Field Plugin (Forster et al., 2004). Correlation of fluorescence and reconstructed electron tomograms was performed using the ec-CLEM Plugin (Paul-Gilloteaux et al., 2017) in ICY (de Chaumont et al., 2012). Alignment was determined by clicking on corresponding pairs of TetraSpeck fiducials in the two imaging modalities.

Tomograms were reconstructed using the IMOD package (Windows Version 6.2) and Etomo (Version 4.9.8, (Kremer et al., 1996)). Patch tracking function was used to perform a fiducial-less image alignment for reconstruction. 3DMOD software was used for manual segmentation of the tomograms. Further editing and annotation were done in Adobe Illustrator (Adobe). Video sequences were compiled in 3DMOD and exported with further editing in ImageJ/FIJI (Schindelin et al., 2012). Video frames were compressed as JPGs to reduce file size.

### 2D electron microscopy

To examine the ultrastructure of *apq12Δ* (CPL1326) and *apq12Δchm7Δ* (CPL1327) strains, unfixed cells were high-pressure frozen using a Leica HMP100 at 2,000 psi and freeze-substituted using a Leica Freeze AFS unit using 1% osmium tetroxide and 1% glutaraldehyde. Samples were infiltrated with durcupan resin (Electron Microscopy Science) and cut in 100 nm thick sections using a Leica UltraCut UC7. Sections were collected on formvar/carbon coated nickel grids and stained with 2% uranyl acetate and lead citrate. Grids were imaged in a FEI Tecnai Biotwin TEM at 80 kV with a Morada CCD camera and iTEM (Olympus) Software.

### Immunogold labeling

For immunogold labeling of nucleoporins, 70 nm Lowicryl sections generated as described above for correlative light and electron tomography were cut using Leica UltraCut UC7 onto 200 mesh copper grids (Quantifoil Micro Tools GmbH). Immunolabeling was carried out with the MAb414 antibody diluted 1:100 in 1% BSA, followed by washes in PBS, and probing with a secondary 10 nm gold-conjugated antibody. After further washes, the grids were fixed in 1% glutaraldehyde in PBS. Lastly, grids were post-stained with 1% uranyl acetate, washed in water and viewed with a Biotwin CM120 Philips equipped with a 1K CCD Camera (Keen View, SIS).

## REFERENCES

Adell MAY, Migliano SM, Upadhyayula S, Bykov YS, Sprenger S, Pakdel M, Vogel GF, Jih G, Skillern W, Behrouzi R, Babst M, Schmidt O, Hess MW, Briggs JA, Kirchhausen T, Teis D. 2017. Recruitment dynamics of ESCRT-III and Vps4 to endosomes and implications for reverse membrane budding. Elife 6:e31652. doi:10.7554/eLife.31652

Adell MAY, Vogel GF, Pakdel M, Müller M, Lindner H, Hess MW, Teis D. 2014. Coordinated binding of Vps4 to ESCRT-III drives membrane neck constriction during MVB vesicle formation. J Cell Biol 205:33–49. doi:10.1083/jcb.201310114

Agromayor M, Carlton JG, Phelan JP, Matthews DR, Carlin LM, Ameer-Beg S, Bowers K, Martin-Serrano J. 2009. Essential role of hIST1 in cytokinesis. Mol Biol Cell 20:1374–87. doi:10.1091/mbc.e08-05-0474

Amberg DC, Burke D, Strathern JN. 2005. Methods in yeast genetics : a Cold Spring Harbor Laboratory course manual.

Arii J, Watanabe M, Maeda F, Tokai-Nishizumi N, Chihara T, Miura M, Maruzuru Y, Koyanagi N, Kato A, Kawaguchi Y. 2018. ESCRT-III mediates budding across the inner nuclear membrane and regulates its integrity. Nat Commun 9:3379. doi:10.1038/s41467-018-05889-9

Bajorek M, Schubert HL, McCullough J, Langelier C, Eckert DM, Stubblefield W-MB, Uter NT, Myszka DG, Hill CP, Sundquist WI. 2009. Structural basis for ESCRT-III protein autoinhibition. Nat Struct Mol Biol 16:754–62. doi:10.1038/nsmb.1621

Barton LJ, Soshnev AA, Geyer PK. 2015. Networking in the nucleus: a spotlight on LEM-domain proteins. Curr Opin Cell Biol 34:1–8. doi:10.1016/j.ceb.2015.03.005

Bauer I, Brune T, Preiss R, Kölling R. 2015. Evidence for a Nonendosomal Function of the Saccharomyces cerevisiae ESCRT-III-Like Protein Chm7. Genetics 201:1439–52. doi:10.1534/genetics.115.178939

Braun DA, Lovric S, Schapiro D, Schneider R, Marquez J, Asif M, Hussain MS, Daga A, Widmeier E, Rao J, Ashraf S, Tan W, Lusk CP, Kolb A, Jobst-Schwan T, Schmidt JM, Hoogstraten CA, Eddy K, Kitzler TM, Shril S, Moawia A, Schrage K, Khayyat AIA, Lawson JA, Gee HY, Warejko JK, Hermle T, Majmundar AJ, Hugo H, Budde B, Motameny S, Altmüller J, Noegel AA, Fathy HM, Gale DP, Waseem SS, Khan A, Kerecuk L, Hashmi S, Mohebbi N, Ettenger R, Serdaroğlu E, Alhasan KA, Hashem M, Goncalves S, Ariceta G, Ubetagoyena M, Antonin W, Baig SM, Alkuraya FS, Shen Q, Xu H, Antignac C, Lifton RP, Mane S, Nürnberg P, Khokha MK, Hildebrandt F. 2018. Mutations in multiple components of the nuclear pore complex cause nephrotic syndrome. J Clin Invest 128:4313–28. doi:10.1172/JCI98688

Braun DA, Sadowski CE, Kohl S, Lovric S, Astrinidis SA, Pabst WL, Gee HY, Ashraf S, Lawson JA, Shril S, Airik M, Tan W, Schapiro D, Rao J, Choi W-I, Hermle T, Kemper MJ, Pohl M, Ozaltin F, Konrad M, Bogdanovic R, Büscher R, Helmchen U, Serdaroglu E, Lifton RP, Antonin W, Hildebrandt F. 2016. Mutations in nuclear pore genes NUP93, NUP205 and XPO5 cause steroid-resistant nephrotic syndrome. Nat Genet 48:457–65. doi:10.1038/ng.3512

Buono RA, Leier A, Paez-Valencia J, Pennington J, Goodman K, Miller N, Ahlquist P, Marquez-Lago TT, Otegui MS. 2017. ESCRT-mediated vesicle concatenation in plant endosomes. J Cell Biol 216:2167–2177. doi:10.1083/jcb.201612040

Caputo S, Couprie J, Duband-Goulet I, Kondé E, Lin F, Braud S, Gondry M, Gilquin B, Worman HJ, Zinn-Justin S. 2006. The carboxyl-terminal nucleoplasmic region of MAN1 exhibits a DNA binding winged helix domain. J Biol Chem 281:18208–15. doi:10.1074/jbc.M601980200

Cashikar AG, Shim S, Roth R, Maldazys MR, Heuser JE, Hanson PI. 2014. Structure of cellular ESCRT-III spirals and their relationship to HIV budding. Elife 3. doi:10.7554/eLife.02184

Christ L, Wenzel EM, Liestøl K, Raiborg C, Campsteijn C, Stenmark H. 2016. ALIX and ESCRT-I/II function as parallel ESCRT-III recruiters in cytokinetic abscission. J Cell Biol 212:499–513. doi:10.1083/jcb.201507009

Colombi P, Webster BM, Fröhlich F, Lusk CP. 2013. The transmission of nuclear pore complexes to daughter cells requires a cytoplasmic pool of Nsp1. J Cell Biol 203:215–32. doi:10.1083/jcb.201305115

Craig R, Beavis RC. 2003. A method for reducing the time required to match protein sequences with tandem mass spectra. Rapid Commun Mass Spectrom 17:2310–16. doi:10.1002/rcm.1198

Cristea IM, Williams R, Chait BT, Rout MP. 2005. Fluorescent proteins as proteomic probes. Mol Cell Proteomics 4:1933–41. doi:10.1074/mcp.M500227-MCP200

Curwin AJ, Brouwers N, Alonso Y Adell M, Teis D, Turacchio G, Parashuraman S, Ronchi P, Malhotra V. 2016. ESCRT-III drives the final stages of CUPS maturation for unconventional protein secretion. Elife 5:e16299. doi:10.7554/eLife.16299

D’Angelo MA, Raices M, Panowski SH, Hetzer MW. 2009. Age-Dependent Deterioration of Nuclear Pore Complexes Causes a Loss of Nuclear Integrity in Postmitotic Cells. Cell 136:284–295. doi:10.1016/j.cell.2008.11.037

de Chaumont F, Dallongeville S, Chenouard N, Hervé N, Pop S, Provoost T, Meas-Yedid V, Pankajakshan P, Lecomte T, Le Montagner Y, Lagache T, Dufour A, Olivo-Marin J-C. 2012. Icy: an open bioimage informatics platform for extended reproducible research. Nat Methods 9:690–6. doi:10.1038/nmeth.2075

de Vos WH, Houben F, Kamps M, Malhas A, Verheyen F, Cox J, Manders EMM, Verstraeten VLRM, Van steensel MAM, Marcelis CLM, Van den wijngaard A, Vaux DJ, Ramaekers FCS, Broers JLV. 2011. Repetitive disruptions of the nuclear envelope invoke temporary loss of cellular compartmentalization in laminopathies. Hum Mol Genet 20:4175–4186. doi:10.1093/hmg/ddr344

Denais CM, Gilbert RM, Isermann P, McGregor AL, te Lindert M, Weigelin B, Davidson PM, Friedl P, Wolf K, Lammerding J. 2016. Nuclear envelope rupture and repair during cancer cell migration. Science 352:353–8. doi:10.1126/science.aad7297

Dou Z, Xu C, Donahue G, Shimi T, Pan JA, Zhu J, Ivanov A, Capell BC, Drake AM, Shah PP, Catanzaro JM, Ricketts MD, Lamark T, Adam SA, Marmorstein R, Zong WX, Johansen T, Goldman RD, Adams PD, Berger SL. 2015. Autophagy mediates degradation of nuclear lamina. Nature 527:105–109. doi:10.1038/nature15548

Doucet CM, Talamas JA, Hetzer MW. 2010. Cell cycle-dependent differences in nuclear pore complex assembly in metazoa. Cell 141:1030–1041. doi:10.1016/j.cell.2010.04.036

Dultz E, Ellenberg J. 2010. Live imaging of single nuclear pores reveals unique assembly kinetics and mechanism in interphase. J Cell Biol 191:15–22. doi:10.1083/jcb.201007076

Fernandez-Martinez J, Phillips J, Sekedat MD, Diaz-Avalos R, Velazquez-Muriel J, Franke JD, Williams R, Stokes DL, Chait BT, Sali A, Rout MP. 2012. Structure-function mapping of a heptameric module in the nuclear pore complex. J Cell Biol 196:419–434. doi:10.1083/jcb.201109008

Floch AG, Palancade B, Doye V. 2014. Fifty years of nuclear pores and nucleocytoplasmic transport studies: multiple tools revealing complex rules. Methods Cell Biol 122:1–40. doi:10.1016/B978-0-12-417160-2.00001-1

Forster B, Van De Ville D, Berent J, Sage D, Unser M. 2004. Complex wavelets for extended depth-of-field: A new method for the fusion of multichannel microscopy images. Microsc Res Tech 65:33–42. doi:10.1002/jemt.20092

Frankel EB, Shankar R, Moresco JJ, Yates JR, Volkmann N, Audhya A. 2017. Ist1 regulates ESCRT-III assembly and function during multivesicular endosome biogenesis in Caenorhabditis elegans embryos. Nat Commun 8:1439. doi:10.1038/s41467-017-01636-8

Freibaum BD, Lu Y, Lopez-Gonzalez R, Kim NC, Almeida S, Lee K-H, Badders N, Valentine M, Miller BL, Wong PC, Petrucelli L, Kim HJ, Gao F-B, Taylor JP. 2015. GGGGCC repeat expansion in C9orf72 compromises nucleocytoplasmic transport. Nature 525:129–33. doi:10.1038/nature14974

Gong Y-N, Guy C, Olauson H, Becker JU, Yang M, Fitzgerald P, Linkermann A, Green DR. 2017. ESCRT-III Acts Downstream of MLKL to Regulate Necroptotic Cell Death and Its Consequences. Cell 169:286–300. doi:10.1016/j.cell.2017.03.020

Gonzalez Y, Saito A, Sazer S. 2012. Fission yeast Lem2 and Man1 perform fundamental functions of the animal cell nuclear lamina. Nucleus 3:60–76. doi:10.4161/nucl.18824

Goodchild RE, Kim CE, Dauer WT. 2005. Loss of the dystonia-associated protein torsinA selectively disrupts the neuronal nuclear envelope. Neuron 48:923–932. doi:10.1016/j.neuron.2005.11.010

Grund SE, Fischer T, Cabal GG, Antúnez O, Pérez-Ortín JE, Hurt E. 2008. The inner nuclear membrane protein Src1 associates with subtelomeric genes and alters their regulated gene expression. J Cell Biol 182:897–910. doi:10.1083/jcb.200803098

Gu M, LaJoie D, Chen OS, von Appen A, Ladinsky MS, Redd MJ, Nikolova L, Bjorkman PJ, Sundquist WI, Ullman KS, Frost A. 2017. LEM2 recruits CHMP7 for ESCRT-mediated nuclear envelope closure in fission yeast and human cells. Proc Natl Acad Sci 114:E2166–E2175. doi:10.1073/pnas.1613916114

Han H, Monroe N, Sundquist WI, Shen PS, Hill CP. 2017. The AAA ATPase Vps4 binds ESCRT-III substrates through a repeating array of dipeptide-binding pockets. Elife 6:e31324. doi:10.7554/eLife.31324

Han H, Monroe N, Votteler J, Shakya B, Sundquist WI, Hill CP. 2015. Binding of Substrates to the Central Pore of the Vps4 ATPase Is Autoinhibited by the Microtubule Interacting and Trafficking (MIT) Domain and Activated by MIT Interacting Motifs (MIMs). J Biol Chem 290:13490–9. doi:10.1074/jbc.M115.642355

Hanson PI, Roth R, Lin Y, Heuser JE. 2008. Plasma membrane deformation by circular arrays of ESCRT-III protein filaments. J Cell Biol 180:389–402. doi:10.1083/jcb.200707031

Hart T, Chandrashekhar M, Aregger M, Steinhart Z, Brown KR, MacLeod G, Mis M, Zimmermann M, Fradet-Turcotte A, Sun S, Mero P, Dirks P, Sidhu S, Roth FP, Rissland OS, Durocher D, Angers S, Moffat J. 2015. High-Resolution CRISPR Screens Reveal Fitness Genes and Genotype-Specific Cancer Liabilities. Cell 163:1515–26. doi:10.1016/j.cell.2015.11.015

Hatch E, Hetzer M. 2014. Breaching the nuclear envelope in development and disease. J Cell Biol 205:133–41. doi:10.1083/jcb.201402003

Hatch EM, Fischer AH, Deerinck TJ, Hetzer MW. 2013. Catastrophic Nuclear Envelope Collapse in Cancer Cell Micronuclei. Cell 154:47–60. doi:10.1016/j.cell.2013.06.007

Hatch EM, Hetzer MW. 2016. Nuclear envelope rupture is induced by actin-based nucleus confinement. J Cell Biol 215:27–36. doi:10.1083/jcb.201603053

Henne WM, Buchkovich NJ, Zhao Y, Emr SD. 2012. The endosomal sorting complex ESCRT-II mediates the assembly and architecture of ESCRT-III helices. Cell 151:356–71. doi:10.1016/j.cell.2012.08.039

Jackson CE, Scruggs BS, Schaffer JE, Hanson PI. 2017. Effects of Inhibiting VPS4 Support a General Role for ESCRTs in Extracellular Vesicle Biogenesis. Biophys J 113:1342–52. doi:10.1016/j.bpj.2017.05.032

Janssens GE, Meinema AC, González J, Wolters JC, Schmidt A, Guryev V, Bischoff R, Wit EC, Veenhoff LM, Heinemann M. 2015. Protein biogenesis machinery is a driver of replicative aging in yeast. Elife 4. doi:10.7554/eLife.08527

Jevtić P, Edens LJ, Vuković LD, Levy DL. 2014. Sizing and shaping the nucleus: mechanisms and significance. Curr Opin Cell Biol 28:16–27. doi:10.1016/j.ceb.2014.01.003

Jimenez AJ, Maiuri P, Lafaurie-Janvore J, Divoux S, Piel M, Perez F. 2014. ESCRT machinery is required for plasma membrane repair. Science 343:1247136. doi:10.1126/science.1247136

Jokhi V, Ashley J, Nunnari J, Noma A, Ito N, Wakabayashi-Ito N, Moore MJ, Budnik V. 2013. Torsin Mediates Primary Envelopment of Large Ribonucleoprotein Granules at the Nuclear Envelope. Cell Rep 3:988–995. doi:10.1016/j.celrep.2013.03.015

Jovičić A, Mertens J, Boeynaems S, Bogaert E, Chai N, Yamada SB, Paul JW, Sun S, Herdy JR, Bieri G, Kramer NJ, Gage FH, Van Den Bosch L, Robberecht W, Gitler AD. 2015. Modifiers of C9orf72 dipeptide repeat toxicity connect nucleocytoplasmic transport defects to FTD/ALS. Nat Neurosci 18:1226–9. doi:10.1038/nn.4085

Kaneb HM, Folkmann AW, Belzil V V, Jao L-E, Leblond CS, Girard SL, Daoud H, Noreau A, Rochefort D, Hince P, Szuto A, Levert A, Vidal S, André-Guimont C, Camu W, Bouchard J-P, Dupré N, Rouleau GA, Wente SR, Dion PA. 2015. Deleterious mutations in the essential mRNA metabolism factor, hGle1, in amyotrophic lateral sclerosis. Hum Mol Genet 24:1363–73. doi:10.1093/hmg/ddu545

Keller A, Nesvizhskii AI, Kolker E, Aebersold R. 2002. Empirical statistical model to estimate the accuracy of peptide identifications made by MS/MS and database search. Anal Chem 74:5383–92.

Kim HJ, Taylor JP. 2017. Lost in Transportation: Nucleocytoplasmic Transport Defects in ALS and Other Neurodegenerative Diseases. Neuron 96:285–97. doi:10.1016/j.neuron.2017.07.029

Kim SJ, Fernandez-Martinez J, Nudelman I, Shi Y, Zhang W, Raveh B, Herricks T, Slaughter BD, Hogan JA, Upla P, Chemmama IE, Pellarin R, Echeverria I, Shivaraju M, Chaudhury AS, Wang J, Williams R, Unruh JR, Greenberg CH, Jacobs EY, Yu Z, de la Cruz MJ, Mironska R, Stokes DL, Aitchison JD, Jarrold MF, Gerton JL, Ludtke SJ, Akey CW, Chait BT, Sali A, Rout MP. 2018. Integrative structure and functional anatomy of a nuclear pore complex. Nature 555:475–482. doi:10.1038/nature26003

King MC, Lusk CP, Blobel G. 2006. Karyopherin-mediated import of integral inner nuclear membrane proteins. Nature 442:1003–1007. doi:10.1038/nature05075

Kosinski J, Mosalaganti S, von Appen A, Teimer R, DiGuilio AL, Wan W, Bui KH, Hagen WJH, Briggs JAG, Glavy JS, Hurt E, Beck M. 2016. Molecular architecture of the inner ring scaffold of the human nuclear pore complex. Science 352:363–5. doi:10.1126/science.aaf0643

Kralt A, Jagalur NB, van den Boom V, Lokareddy RK, Steen A, Cingolani G, Fornerod M, Veenhoff LM. 2015. Conservation of inner nuclear membrane targeting sequences in mammalian Pom121 and yeast Heh2 membrane proteins. Mol Biol Cell 26:3301–3312. doi:10.1091/mbc.E15-03-0184

Kremer JR, Mastronarde DN, McIntosh JR. 1996. Computer visualization of three-dimensional image data using IMOD. J Struct Biol 116:71–6. doi:10.1006/jsbi.1996.0013

Kukulski W, Schorb M, Welsch S, Picco A, Kaksonen M, Briggs JAG. 2012. Precise, Correlated Fluorescence Microscopy and Electron Tomography of Lowicryl Sections Using Fluorescent Fiducial Markers. pp. 235–257. doi:10.1016/B978-0-12-416026-2.00013-3

Kulak NA, Pichler G, Paron I, Nagaraj N, Mann M. 2014. Minimal, encapsulated proteomic-sample processing applied to copy-number estimation in eukaryotic cells. Nat Methods 11:319–24. doi:10.1038/nmeth.2834

LaCava J, Fernandez-Martinez J, Hakhverdyan Z, Rout MP. 2016. Optimized Affinity Capture of Yeast Protein Complexes. Cold Spring Harb Protoc 2016:pdb.prot087932. doi:10.1101/pdb.prot087932

Lata S, Roessle M, Solomons J, Jamin M, Gottlinger HG, Svergun DI, Weissenhorn W. 2008. Structural basis for autoinhibition of ESCRT-III CHMP3. J Mol Biol 378:818–27. doi:10.1016/j.jmb.2008.03.030

Laudermilch E, Tsai P-L, Graham M, Turner E, Zhao C, Schlieker C. 2016. Dissecting Torsin/cofactor function at the nuclear envelope: a genetic study. Mol Biol Cell 27:3964–71. doi:10.1091/mbc.E16-07-0511

Lee C-P, Liu G-T, Kung H-N, Liu P-T, Liao Y-T, Chow L-P, Chang L-S, Chang Y-H, Chang C-W, Shu W-C, Angers A, Farina A, Lin S-F, Tsai C-H, Bouamr F, Chen M-R. 2016. The Ubiquitin Ligase Itch and Ubiquitination Regulate BFRF1-Mediated Nuclear Envelope Modification for Epstein-Barr Virus Maturation. J Virol 90:8994–9007. doi:10.1128/JVI.01235-16

Lee CP, Liu PT, Kung HN, Su MT, Chua HH, Chang YH, Chang CW, Tsai CH, Liu FT, Chen MR. 2012. The ESCRT Machinery Is Recruited by the Viral BFRF1 Protein to the Nucleus-Associated Membrane for the Maturation of Epstein-Barr Virus. PLoS Pathog 8:e1002904. doi:10.1371/journal.ppat.1002904

Li X, Qian J, Wang C, Zheng K, Ye L, Fu Y, Han N, Bian H, Pan J, Wang J, Zhu M. 2011. Regulating Cytoplasmic Calcium Homeostasis Can Reduce Aluminum Toxicity in Yeast. PLoS One 6:e21148. doi:10.1371/journal.pone.0021148

Lokareddy RK, Hapsari RA, Van rheenen M, Pumroy RA, Bhardwaj A, Steen A, Veenhoff LM, Cingolani G. 2015. Distinctive Properties of the Nuclear Localization Signals of Inner Nuclear Membrane Proteins Heh1 and Heh2. Structure 23:1305–1316. doi:10.1016/j.str.2015.04.017

Longtine MS, McKenzie A, Demarini DJ, Shah NG, Wach A, Brachat A, Philippsen P, Pringle JR. 1998. Additional modules for versatile and economical PCR-based gene deletion and modification in Saccharomyces cerevisiae. Yeast 14:953–961. doi:10.1002/

Lord CL, Timney BL, Rout MP, Wente SR. 2015. Altering nuclear pore complex function impacts longevity and mitochondrial function in S. cerevisiae. J Cell Biol 208:729–44. doi:10.1083/jcb.201412024

Lusk CP, Blobel G, King MC. 2007. Highway to the inner nuclear membrane: rules for the road. Nat Rev Mol Cell Biol 8:414–20. doi:10.1038/nrm2165

Lusk CP, King MC. 2017. The nucleus: keeping it together by keeping it apart. Curr Opin Cell Biol 44:44–50. doi:10.1016/j.ceb.2017.02.001

Maciejowski J, Li Y, Bosco N, Campbell PJ, de Lange T. 2015. Chromothripsis and Kataegis Induced by Telomere Crisis. Cell 163:1641–54. doi:10.1016/j.cell.2015.11.054

Makio T, Lapetina DL, Wozniak RW. 2013. Inheritance of yeast nuclear pore complexes requires the Nsp1p subcomplex. J Cell Biol 203:187–96. doi:10.1083/jcb.201304047

Malhas A, Goulbourne C, Vaux DJ. 2011. The nucleoplasmic reticulum: form and function. Trends Cell Biol 21:362–73. doi:10.1016/j.tcb.2011.03.008

Mans BJ, Anantharaman V, Aravind L, Koonin E V, Wang Z, Casciola-Rosen L, Rosen A. 2004. Comparative genomics, evolution and origins of the nuclear envelope and nuclear pore complex. Cell Cycle 3:1612–37. doi:10.4161/cc.3.12.1345

Maul GG, Maul HM, Scogna JE, Lieberman MW, Stein GS, Hsu BY, Borun TW. 1972. Time Sequence of Nuclear Pore Formation in Phytohemagglutinin-Stimulated Lymphocytes and in Hela Cells During the Cell Cycle. J Cell Biol 55:433–447. doi:10.1083/jcb.55.2.433

McCullough J, Clippinger AK, Talledge N, Skowyra ML, Saunders MG, Naismith T V., Colf LA, Afonine P, Arthur C, Sundquist WI, Hanson PI, Frost A. 2015. Structure and membrane remodeling activity of ESCRT-III helical polymers. Science 350:1548–51. doi:10.1126/science.aad8305

McCullough J, Frost A, Sundquist WI. 2018. Structures, Functions, and Dynamics of ESCRT-III/Vps4 Membrane Remodeling and Fission Complexes. Annu Rev Cell Dev Biol 34:85–109. doi:10.1146/annurev-cellbio-100616-060600

Meinema AC, Laba JK, Hapsari RA, Otten R, Mulder FAA, Kralt A, van den Bogaart G, Lusk CP, Poolman B, Veenhoff LM. 2011. Long unfolded linkers facilitate membrane protein import through the nuclear pore complex. Science 333:90–3. doi:10.1126/science.1205741

Mekhail K, Seebacher J, Gygi SP, Moazed D. 2008. Role for perinuclear chromosome tethering in maintenance of genome stability. Nature 456:667–670. doi:10.1038/nature07460

Miyake N, Tsukaguchi H, Koshimizu E, Shono A, Matsunaga S, Shiina M, Mimura Y, Imamura S, Hirose T, Okudela K, Nozu K, Akioka Y, Hattori M, Yoshikawa N, Kitamura A, Cheong H Il, Kagami S, Yamashita M, Fujita A, Miyatake S, Tsurusaki Y, Nakashima M, Saitsu H, Ohashi K, Imamoto N, Ryo A, Ogata K, Iijima K, Matsumoto N. 2015. Biallelic Mutations in Nuclear Pore Complex Subunit NUP107 Cause Early-Childhood-Onset Steroid-Resistant Nephrotic Syndrome. Am J Hum Genet 97:555–66. doi:10.1016/j.ajhg.2015.08.013

Mochida K, Oikawa Y, Kimura Y, Kirisako H, Hirano H, Ohsumi Y, Nakatogawa H. 2015. Receptor-mediated selective autophagy degrades the endoplasmic reticulum and the nucleus. Nature 522:359–362. doi:10.1038/nature14506

Monroe N, Han H, Shen PS, Sundquist WI, Hill CP. 2017. Structural basis of protein translocation by the Vps4-Vta1 AAA ATPase. Elife 6:e24487. doi:10.7554/eLife.24487

Morita E, Sandrin V, McCullough J, Katsuyama A, Baci Hamilton I, Sundquist WI. 2011. ESCRT-III Protein Requirements for HIV-1 Budding. Cell Host Microbe 9:235–42. doi:10.1016/j.chom.2011.02.004

Mostofa MG, Rahman MA, Koike N, Yeasmin AM, Islam N, Waliullah TM, Hosoyamada S, Shimobayashi M, Kobayashi T, Hall MN, Ushimaru T. 2018. CLIP and cohibin separate rDNA from nucleolar proteins destined for degradation by nucleophagy. J Cell Biol 217:2675–90. doi:10.1083/jcb.201706164

Nesvizhskii AI, Keller A, Kolker E, Aebersold R. 2003. A statistical model for identifying proteins by tandem mass spectrometry. Anal Chem 75:4646–58.

Neville M, Rosbash M. 1999. The NES-Crm1p export pathway is not a major mRNA export route in Saccharomyces cerevisiae. EMBO J 18:3746–56. doi:10.1093/emboj/18.13.3746

Nousiainen HO, Kestilä M, Pakkasjärvi N, Honkala H, Kuure S, Tallila J, Vuopala K, Ignatius J, Herva R, Peltonen L. 2008. Mutations in mRNA export mediator GLE1 result in a fetal motoneuron disease. Nat Genet 40:155–7. doi:10.1038/ng.2007.65

Obita T, Saksena S, Ghazi-Tabatabai S, Gill DJ, Perisic O, Emr SD, Williams RL. 2007. Structural basis for selective recognition of ESCRT-III by the AAA ATPase Vps4. Nature 449:735–739. doi:10.1038/nature06171

Oeffinger M, Wei KE, Rogers R, DeGrasse JA, Chait BT, Aitchison JD, Rout MP. 2007. Comprehensive analysis of diverse ribonucleoprotein complexes. Nat Methods 4:951–956. doi:10.1038/nmeth1101

Olmos Y, Hodgson L, Mantell J, Verkade P, Carlton JG. 2015. ESCRT-III controls nuclear envelope reformation. Nature 522:236–239. doi:10.1038/nature14503

Olmos Y, Perdrix-Rosell A, Carlton JG. 2016. Membrane Binding by CHMP7 Coordinates ESCRT-III-Dependent Nuclear Envelope Reformation. Curr Biol 26:2635–41. doi:10.1016/j.cub.2016.07.039

Onischenko E, Tang JH, Andersen KR, Knockenhauer KE, Vallotton P, Derrer CP, Kralt A, Mugler CF, Chan LY, Schwartz TU, Weis K. 2017. Natively Unfolded FG Repeats Stabilize the Structure of the Nuclear Pore Complex. Cell 171:904–17.e19. doi:10.1016/j.cell.2017.09.033

Otsuka S, Bui KH, Schorb M, Hossain MJ, Politi AZ, Koch B, Eltsov M, Beck M, Ellenberg J. 2016. Nuclear pore assembly proceeds by an inside-out extrusion of the nuclear envelope. Elife 5. doi:10.7554/eLife.19071

Pappas SS, Liang C-C, Kim S, Rivera CO, Dauer WT. 2018. TorsinA dysfunction causes persistent neuronal nuclear pore defects. Hum Mol Genet 27:407–420. doi:10.1093/hmg/ddx405

Paul-Gilloteaux P, Heiligenstein X, Belle M, Domart M-C, Larijani B, Collinson L, Raposo G, Salamero J. 2017. eC-CLEM: flexible multidimensional registration software for correlative microscopies. Nat Methods 14:102–3. doi:10.1038/nmeth.4170

Popken P, Ghavami A, Onck PR, Poolman B, Veenhoff LM. 2015. Size-dependent leak of soluble and membrane proteins through the yeast nuclear pore complex. Mol Biol Cell 26:1386–94. doi:10.1091/mbc.E14-07-1175

Raab M, Gentili M, de Belly H, Thiam HR, Vargas P, Jimenez AJ, Lautenschlaeger F, Voituriez R, Lennon-Duménil AM, Manel N, Piel M. 2016. ESCRT III repairs nuclear envelope ruptures during cell migration to limit DNA damage and cell death. Science 352:359–62. doi:10.1126/science.aad7611

Radulovic M, Schink KO, Wenzel EM, Nähse V, Bongiovanni A, Lafont F, Stenmark H. 2018. ESCRT-mediated lysosome repair precedes lysophagy and promotes cell survival. EMBO J 37:e99753. doi:10.15252/embj.201899753

Roberts P, Moshitch-Moshkovitz S, Kvam E, O’toole E, Winey M, Goldfarb DS, O’Toole E, Winey M, Goldfarb DS. 2003. Piecemeal microautophagy of nucleus in Saccharomyces cerevisiae. Mol Biol Cell 14:129–41. doi:10.1091/mbc.E02

Robijns J, Molenberghs F, Sieprath T, Corne TDJ, Verschuuren M, De Vos WH. 2016. In silico synchronization reveals regulators of nuclear ruptures in lamin A/C deficient model cells. Sci Rep 6:30325. doi:10.1038/srep30325

Romanauska A, Köhler A. 2018. The Inner Nuclear Membrane Is a Metabolically Active Territory that Generates Nuclear Lipid Droplets. Cell 174:700–15. doi:10.1016/j.cell.2018.05.047

Saksena S, Wahlman J, Teis D, Johnson AE, Emr SD. 2009. Functional reconstitution of ESCRT-III assembly and disassembly. Cell 136:97–109. doi:10.1016/j.cell.2008.11.013

Savas JN, Toyama BH, Xu T, Yates JR, Hetzer MW. 2012. Extremely long-lived nuclear pore proteins in the rat brain. Science 335:942. doi:10.1126/science.1217421

Scarcelli JJ, Hodge CA, Cole CN. 2007. The yeast integral membrane protein Apq12 potentially links membrane dynamics to assembly of nuclear pore complexes. J Cell Biol 178:799–812. doi:10.1083/jcb.200702120

Scheffer LL, Sreetama SC, Sharma N, Medikayala S, Brown KJ, Defour A, Jaiswal JK. 2014. Mechanism of Ca^2+^-triggered ESCRT assembly and regulation of cell membrane repair. Nat Commun 5:5646. doi:10.1038/ncomms6646

Schindelin J, Arganda-Carreras I, Frise E, Kaynig V, Longair M, Pietzsch T, Preibisch S, Rueden C, Saalfeld S, Schmid B. 2012. Fiji: an open-source platform for biological-image analysis. Nat Methods 9:676–682.

Schmidt HB, Görlich D. 2016. Transport Selectivity of Nuclear Pores, Phase Separation, and Membraneless Organelles. Trends Biochem Sci 41:46–61. doi:10.1016/j.tibs.2015.11.001

Schöneberg J, Lee I-H, Iwasa JH, Hurley JH. 2017. Reverse-topology membrane scission by the ESCRT proteins. Nat Rev Mol Cell Biol 18:5–17. doi:10.1038/nrm.2016.121

Schöneberg J, Pavlin MR, Yan S, Righini M, Lee I-H, Carlson L-A, Bahrami AH, Goldman DH, Ren X, Hummer G, Bustamante C, Hurley JH. 2018. ATP-dependent force generation and membrane scission by ESCRT-III and Vps4. Science 362:1423–8. doi:10.1126/science.aat1839

Schreiber KH, Kennedy BK. 2013. When lamins go bad: nuclear structure and disease. Cell 152:1365–75. doi:10.1016/j.cell.2013.02.015

Schreiner SM, Koo PK, Zhao Y, Mochrie SGJ, King MC. 2015. The tethering of chromatin to the nuclear envelope supports nuclear mechanics. Nat Commun 6:7159. doi:10.1038/ncomms8159

Searle BC, Turner M, Nesvizhskii AI. 2008. Improving Sensitivity by Probabilistically Combining Results from Multiple MS/MS Search Methodologies. J Proteome Res 7:245–53. doi:10.1021/pr070540w

Serebryannyy L, Misteli T. 2018. Protein sequestration at the nuclear periphery as a potential regulatory mechanism in premature aging. J Cell Biol 217:21–37. doi:10.1083/jcb.201706061

Shi KY, Mori E, Nizami ZF, Lin Y, Kato M, Xiang S, Wu LC, Ding M, Yu Y, Gall JG, McKnight SL. 2017. Toxic PRn poly-dipeptides encoded by the C9orf72 repeat expansion block nuclear import and export. Proc Natl Acad Sci U S A 114:E1111–E1117. doi:10.1073/pnas.1620293114

Shim S, Kimpler LA, Hanson PI. 2007. Structure/Function Analysis of Four Core ESCRT-III Proteins Reveals Common Regulatory Role for Extreme C-Terminal Domain. Traffic 8:1068–1079. doi:10.1111/j.1600-0854.2007.00584.x

Skowyra ML, Schlesinger PH, Naismith T V., Hanson PI. 2018. Triggered recruitment of ESCRT machinery promotes endolysosomal repair. Science 360:5078. doi:10.1126/science.aar5078

Speese SD, Ashley J, Jokhi V, Nunnari J, Barria R, Li Y, Ataman B, Koon A, Chang Y-T, Li Q, Moore MJ, Budnik V. 2012. Nuclear Envelope Budding Enables Large Ribonucleoprotein Particle Export during Synaptic Wnt Signaling. Cell 149:832–846. doi:10.1016/j.cell.2012.03.032

Stuchell-Brereton MD, Skalicky JJ, Kieffer C, Karren MA, Ghaffarian S, Sundquist WI. 2007. ESCRT-III recognition by VPS4 ATPases. Nature 449:740–4. doi:10.1038/nature06172

Su M, Guo EZ, Ding X, Li Y, Tarrasch JT, Brooks CL, Xu Z, Skiniotis G. 2017. Mechanism of Vps4 hexamer function revealed by cryo-EM. Sci Adv 3:e1700325. doi:10.1126/sciadv.1700325

Tang S, Buchkovich NJ, Henne WM, Banjade S, Kim YJ, Emr SD. 2016. ESCRT-III activation by parallel action of ESCRT-I/II and ESCRT-0/Bro1 during MVB biogenesis. Elife 5:e15507. doi:10.7554/eLife.15507

Tang S, Henne WM, Borbat PP, Buchkovich NJ, Freed JH, Mao Y, Fromme JC, Emr SD. 2015. Structural basis for activation, assembly and membrane binding of ESCRT-III Snf7 filaments. Elife 4:e12548. doi:10.7554/eLife.12548

Teis D, Saksena S, Judson BL, Emr SD. 2010. ESCRT-II coordinates the assembly of ESCRT-III filaments for cargo sorting and multivesicular body vesicle formation. EMBO J 29:871–83. doi:10.1038/emboj.2009.408

Thaller DJ, Lusk CP. 2018. Fantastic nuclear envelope herniations and where to find them. Biochem Soc Trans 46:877–89. doi:10.1042/BST20170442

Timney BL, Raveh B, Mironska R, Trivedi JM, Kim SJ, Russel D, Wente SR, Sali A, Rout MP. 2016. Simple rules for passive diffusion through the nuclear pore complex. J Cell Biol 215:57–76. doi:10.1083/jcb.201601004

Toyama BH, Arrojo E Drigo R, Lev-Ram V, Ramachandra R, Deerinck TJ, Lechene C, Ellisman MH, Hetzer MW. 2018. Visualization of long-lived proteins reveals age mosaicism within nuclei of postmitotic cells. J Cell Biol 1–16. doi:10.1083/jcb.201809123

Toyama BH, Savas JN, Park SK, Harris MS, Ingolia NT, Yates JR, Hetzer MW. 2013. Identification of long-lived proteins reveals exceptional stability of essential cellular structures. Cell 154:971–82. doi:10.1016/j.cell.2013.07.037

Van Driessche B, Tafforeau L, Hentges P, Carr AM, Vandenhaute J. 2005. Additional vectors for PCR-based gene tagging in Saccharomyces cerevisiae and Schizosaccharomyces pombe using nourseothricin resistance. Yeast 22:1061–8. doi:10.1002/yea.1293

Vargas JD, Hatch EM, Anderson DJ, Hetzer MW. 2012. Transient nuclear envelope rupturing during interphase in human cancer cells. Nucleus 3:88–100. doi:10.4161/nucl.18954

Ventimiglia LN, Cuesta-Geijo MA, Martinelli N, Caballe A, Macheboeuf P, Miguet N, Parnham IM, Olmos Y, Carlton JG, Weissenhorn W, Martin-Serrano J. 2018. CC2D1B Coordinates ESCRT-III Activity during the Mitotic Reformation of the Nuclear Envelope. Dev Cell 47:547–63.e6. doi:10.1016/j.devcel.2018.11.012

Vietri M, Schink KO, Campsteijn C, Wegner CS, Schultz SW, Christ L, Thoresen SB, Brech A, Raiborg C, Stenmark H. 2015. Spastin and ESCRT-III coordinate mitotic spindle disassembly and nuclear envelope sealing. Nature 522:231–5. doi:10.1038/nature14408

von Schwedler UK, Stuchell M, Müller B, Ward DM, Chung H-Y, Morita E, Wang HE, Davis T, He G-P, Cimbora DM, Scott A, Kräusslich H-G, Kaplan J, Morham SG, Sundquist WI. 2003. The protein network of HIV budding. Cell 114:701–13.

Webster BM, Colombi P, Jäger J, Lusk CP. 2014. Surveillance of nuclear pore complex assembly by ESCRT-III/Vps4. Cell 159:388–401. doi:10.1016/j.cell.2014.09.012

Webster BM, Thaller DJ, Jäger J, Ochmann SE, Borah S, Lusk CP. 2016. Chm7 and Heh1 collaborate to link nuclear pore complex quality control with nuclear envelope sealing. EMBO J 35:2447–67. doi:10.15252/embj.201694574

Wemmer M, Azmi I, West M, Davies B, Katzmann D, Odorizzi G. 2011. Bro1 binding to Snf7 regulates ESCRT-III membrane scission activity in yeast. J Cell Biol 192:295–306. doi:10.1083/jcb.201007018

Wente SR, Blobel G. 1993. A temperature-sensitive NUP116 null mutant forms a nuclear envelope seal over the yeast nuclear pore complex thereby blocking nucleocytoplasmic traffic. J Cell Biol 123:275–84. doi:10.1083/jcb.123.2.275

Wenzel EM, Schultz SW, Schink KO, Pedersen NM, Nähse V, Carlson A, Brech A, Stenmark H, Raiborg C. 2018. Concerted ESCRT and clathrin recruitment waves define the timing and morphology of intraluminal vesicle formation. Nat Commun 9:2932. doi:10.1038/s41467-018-05345-8

West M, Zurek N, Hoenger A, Voeltz GK. 2011. A 3D analysis of yeast ER structure reveals how ER domains are organized by membrane curvature. J Cell Biol 193:333–346. doi:10.1083/jcb.201011039

Winey M, Yarar D, Giddings TH, Mastronarde DN. 1997. Nuclear pore complex number and distribution throughout the Saccharomyces cerevisiae cell cycle by three-dimensional reconstruction from electron micrographs of nuclear envelopes. Mol Biol Cell 8:2119–32. doi:10.1091/mbc.8.11.2119

Xiao J, Chen X-W, Davies BA, Saltiel AR, Katzmann DJ, Xu Z. 2009. Structural basis of Ist1 function and Ist1-Did2 interaction in the multivesicular body pathway and cytokinesis. Mol Biol Cell 20:3514–24. doi:10.1091/mbc.e09-05-0403

Xu D, Marquis K, Pei J, Fu S-C, Cağatay T, Grishin N V, Chook YM. 2015. LocNES: a computational tool for locating classical NESs in CRM1 cargo proteins. Bioinformatics 31:1357–65. doi:10.1093/bioinformatics/btu826

Yam C, Gu Y, Oliferenko S. 2013. Partitioning and remodeling of the Schizosaccharomyces japonicus mitotic nucleus require chromosome tethers. Curr Biol 23:2303–10. doi:10.1016/j.cub.2013.09.057

Yang B, Stjepanovic G, Shen Q, Martin A, Hurley JH. 2015. Vps4 disassembles an ESCRT-III filament by global unfolding and processive translocation. Nat Struct Mol Biol 22:492–8. doi:10.1038/nsmb.3015

Yewdell WT, Colombi P, Makhnevych T, Lusk CP. 2011. Lumenal interactions in nuclear pore complex assembly and stability. Mol Biol Cell 22:1375–88. doi:10.1091/mbc.E10-06-0554

Zamborlini A, Usami Y, Radoshitzky SR, Popova E, Palu G, Gottlinger H. 2006. Release of autoinhibition converts ESCRT-III components into potent inhibitors of HIV-1 budding. Proc Natl Acad Sci 103:19140–5. doi:10.1073/pnas.0603788103

Zhang K, Donnelly CJ, Haeusler AR, Grima JC, Machamer JB, Steinwald P, Daley EL, Miller SJ, Cunningham KM, Vidensky S, Gupta S, Thomas MA, Hong I, Chiu S-L, Huganir RL, Ostrow LW, Matunis MJ, Wang J, Sattler R, Lloyd TE, Rothstein JD. 2015. The C9orf72 repeat expansion disrupts nucleocytoplasmic transport. Nature 525:56–61. doi:10.1038/nature14973

Zhang W, Neuner A, Rüthnick D, Sachsenheimer T, Lüchtenborg C, Brügger B, Schiebel E. 2018. Brr6 and Brl1 locate to nuclear pore complex assembly sites to promote their biogenesis. J Cell Biol 217:877–94. doi:10.1083/jcb.201706024

